# Single *MAPT* knock-in mouse models of frontotemporal dementia for sharing with the neurodegenerative research community

**DOI:** 10.1101/2024.11.15.623736

**Authors:** Takahiro Morito, Mohan Qi, Naoko Kamano, Hiroki Sasaguri, Sumi Bez, Martha Foiani, Keren Duff, Seico Benner, Toshihiro Endo, Hiroshi Hama, Hiroshi Kurokawa, Atushi Miyawaki, Hiroshi Mizuma, Naruhiko Sahara, Masafumi Shimojo, Makoto Higuchi, Takaomi C. Saido, Naoto Watamura

## Abstract

We recently reported development of human *MAPT* knock-in mice that carry single or double pathogenic mutations of frontotemporal dementia. However, it takes more than 14 months for the line with the most aggressive phenotypes to exhibit tau pathology without forming high-order tau oligomers, along with concomitant abnormal behavior. We thus generated *MAPT* knock-in mice carrying triple mutations, among which the *MAPT*^P301S;Int10+3;S320F^ line exhibited robust pathology starting earlier than 6 months. Tau accumulation took place mainly in the thalamus, hypothalamus, amygdala and entorhinal cortex, but less so in the hippocampus, leading to synaptic loss, atrophy and behavioral abnormalities. Crossbreeding *MAPT*^P301S;Int10+3;S320F^ with *App* knock-in mice *App*^NL-G-F^ resulted in the manifestation of tau pathology in the hippocampus and cortex. These mutant mice will be valuable tools for understanding the mechanisms of frontotemporal dementia, Alzheimer’s disease and other tauopathies.

## Introduction

Mutations in the coding and non-coding regions of the human *MAPT* gene are known to cause a form of frontotemporal dementia (FTD) ^1^. Accumulation of hyperphosphorylated tau in the brain represents the major pathological hallmark of FTD and Parkinsonism linked to chromosome 17 (FTDP-17). We previously introduced FTD-causing pathogenic mutations into human *MAPT* knock-in (KI) mice ^2,3^ using cytosine base editor (BE) ^4^, which resulted in the generation of seven isogenic mutant KI mouse lines^5^ (**Table 1**). Utilizing KI mice rather than transgenic overexpression mice is crucial for modeling diseased conditions while avoiding overexpression artifacts such as the destruction of endogenous loci, disturbances in cellular protein localization/interactions, and non-specific endoplasmic reticulum stress/calcium mobilization ^6–12^. Another major weakness of the overexpression paradigm is the inapplicability of genome editing because multiple copies of cDNA(s) are usually inserted into the host genome in a random manner ^13^.

**Table 1.** Mutant human *MAPT* mouse lines generated. Lines 8-10 represent newly generated lines in the present study.

| Lines | Mutation(s) | References | AT8 positive tau | Off-target analysis |
| --- | --- | --- | --- | --- |
| 1 | P301L | Watamura et al. (2024) | N/A | No |
| 2 | P301S | Watamura et al. (2024) | N/A | No |
| 3 | P301V | Watamura et al. (2024) | N/A | No |
| 4 | Int10+3 G>A | Watamura et al. (2024) | After 14 months | Yes |
| 5 | P301L; Int10+3 G>A | Watamura et al. (2024) | After 14 months | No |
| 6 | P301S; Int10+3 G>A | Watamura et al. (2024); Saido (2024) | After 24 months | Yes |
| 7 | S305N; Int10+3 G>A | Watamura et al. (2024) | After 12 months | Yes |
| 8 | P301S; Int10+3 G>A; S320F | This paper | Before 6 months | Yes |
| 9 | S305N; Int10+3 G>A; S320F | This paper | After 9 months | Yes |
| 10 | S305N; Int10+3 G>A; S320Y | This paper | Around 6 months | Yes |
| 11 | S320F | N/A | N/A | No |
| 12 | Int10+3 G>A, S320F | N/A | N/A | No |
| 13 | P301L; S320F | N/A | N/A | No |

Of the seven lines mentioned above, we had made detailed immunochemical and biochemical analyses of two KI lines that showed prominent tau pathology: a single Intron10+3 G>A mutant and a double S305N-Intron10+3 G>A mutant (*MAPT*^Int10+3^ and *MAPT*^S305N;Int10+3^). Both these lines exhibited alterations in *MAPT* pre-mRNA splicing and a significant increase in 4-repeat (4R) tau isoforms with a concomitant decrease in 3-repeat (3R) isoforms, resulting in the hyperphosphorylation of tau protein ^5^. These mutant mice presented pathological deposition of hyperphosphorylated tau accompanying synaptic loss and cognitive impairment at the age of 15 months or later, but they failed to show accumulation of seed-competent tau oligomers as in human cases. Moreover, they are not user-friendly models because the pathological and behavioral changes they manifest proceed relatively slowly and only to a mild extent. We also created another double mutant *MAPT* KI mouse, *MAPT*^P301S;Int10+3^, but this line also showed almost no phenotype until their 24 months of age unless crossbred with *App* knock-in mice ^11^.

We previously showed that a triple mutant *App* KI line, *App*^NL-G-F^, exhibits much earlier (approximately 3 times) and more aggressive Aβ pathology than the double mutant line, *App*^NL-F 6,11^, demonstrating that a combination of multiple mutations can synergistically strengthen the pathological phenotypes in mice. To generate pathophysiologically more relevant and user-friendly models of FTD, we therefore added another pathogenic mutation, S320F, to *MAPT*^S305N; Int10+3^ and *MAPT* ^P301S;Int10+3^ KI mice. This mutation, present in exon 11 of the *MAPT* gene, is known to affect the structural properties of tau protein rather than the alternative splicing of its pre-mRNA ^14–17^. These efforts resulted in the generation of the new triple mutant lines, i.e. *MAPT*^P301S;Int10+3;S320F^, *MAPT*^S305N;Int10+3;S320F^ and *MAPT*^S305N;Int10+3;S320Y^ (Table 1).

We herein describe the pathological, structural, biochemical and behavioral phenotypes of the triple mutant KI lines. Among the three, the *MAPT* ^P301S;Int10+3;S320F^ line exhibited typical FTD-like pathology accompanying hyperphosphorylated oligomeric tau formation together with synaptic loss earlier than 6 months of age. We also observed local brain atrophy, FTD-like behavioral abnormality and an increase in body weight at a later age (9-12 months). Accordingly, these next-generation mouse models of FTD should be suitable for use in preclinical studies to understand the disease mechanisms and to search for therapeutic candidates not only for FTD but also for Alzheimer’s disease (AD), with minimal concerns about overexpression artifacts in both *in vivo* and *in vitro* contexts.

## Results

### Generation of mutant *MAPT* KI mice that harbor three FTD-causing mutations

We previously created two mouse lines that harbor S305N/Intron10+3 G>C mutations (*MAPT*^S305N;Int10+3^) or P301S/Intron10+3 G>C mutations (*MAPT*^P301S;Int10+3^) using cytosine BE ^4^, but the progression of their pathology took over a year to become apparent (**Table 1**) ^5,11^. Because an S320F mutation have been reported to accelerate aggregation of tau protein in combination with P301 mutations ^14,17^, we added this mutation to the double mutants by designing a sgRNA targeting the S320 position with -NGG PAM sequences for BE-mediated genome editing. The sgRNA and cytosine BE mRNA were microinjected into the zygotes from *MAPT*^S305N;Int10+3^ and *MAPT*^P301S;Int10+3^ mice, and the zygotes were individualized to obtain triple mutant mice. We successfully introduced an S320F mutation to *MAPT*^S305N;Int10+3^ and *MAPT*^P301S;Int10+3^ KI mice in the second-round genome editing, thereby obtaining the following lines: *MAPT*^S305N;Int10+3^ mice with the S320F mutation (*MAPT*^S305N;Int10+3;S320F^) and *MAPT*^P301S;Int10+3^ mice with the S320F mutation (*MAPT*^P301S;Int10+3;S320F^) (**Figure 1A-C**) (**Table 1**). After the second editing, we confirmed the absence of any potential off-target mutations by *in silico* homology analysis and by whole genome resequencing analysis (**Supplementary Table 1**) ^18^. After backcrossing the mutants with wild-type B6/J mice for three times or more, we obtained homozygous *MAPT*^S305N;Int10+3;S320F^ and *MAPT*^P301S;Int10+3;S320F^. All the mice described in this paper are homozygotes. Due presumably to the random nature of BE, we also obtained *MAPT*^S305N;Int10+3^ mice carrying the S320Y mutation (*MAPT*^S305N;Int10+3;S320Y^) (**Table 1**). The other mutants generated during these procedures include the following: the double *MAPT*-Intron10+3 G>A plus S320F mutation (*MAPT*^Int10+3;S320F^), *MAPT*-P301L plus S320F mutation (*MAPT*^P301L;S320F^) and the single *MAPT*-S320F mutation (*MAPT*^S320F^) (**Supplementary Figure 1**, **Table 1**).

**Figure 1.**
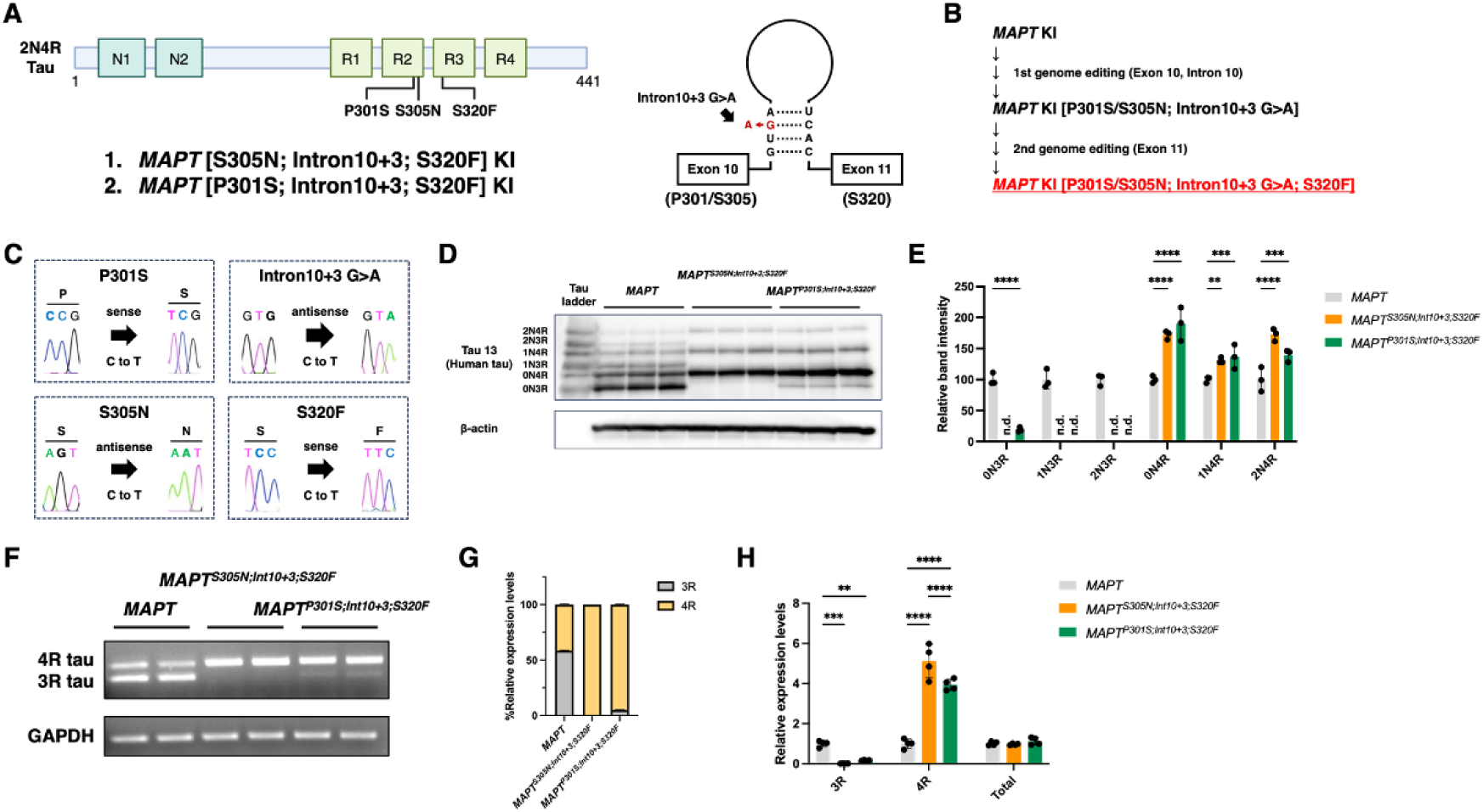
Generation and isoform expression patterns of triple mutant *MAPT* KI mouse lines. **A.** Pathogenic mutations of human *MAPT* gene in the exon (left) and intron (right). **B.** Workflow of the two genome editing steps for generating *MAPT*^S305N;Int10+3;S320F^ and *MAPT*^P301S;Int10+3;S320F^ KI mice. **C.** Four mutations introduced via cytosine base editing. **D, E.** Western blot analysis of alkaline phosphatase-treated tau protein from the *MAPT*^S305N;Int10+3;S320F^ and *MAPT*^P301S;Int10+3;S320F^ KI mouse frontal cortex (D) and densitometric quantification of each isoform (E). **F, G.** RT-PCR analysis of the *MAPT* gene transcripts using 3R-and 4R-specific primer sets (F) and densitometric quantification (G). **H.** Quantitative PCR analysis of the *MAPT* gene transcripts. Statistical significance was examined by two-way ANOVA (E, H). **: p<0.01, ***: p<0.005, ****: p<0.001. n.d.: not detected.

### Tau isoforms of triple mutant *MAPT* KI mice

To investigate the expression pattern of tau isoforms, we performed Western blot analysis using brain homogenates from cortices of each mouse line after dephosphorylation treatment (**Figure 1D**). The expression of 3R tau was undetectable in *MAPT*^S305N;Int10+3;S320F^ mice. In contrast, the *MAPT*^P301S;Int10+3;S320F^ line produced a small but detectable quantity of 3R tau, the 0N3R isoform in particular (**Figure 1E**). Accordingly, the 4R tau was highly expressed in the *MAPT*^S305N;Int10+3;S320F^ mice, and the MAPT^P301S;Int10+3;S320F^ _mice expressed much_ more 4R than 3R tau isoforms (**Figure 1E**). These results are consistent with the observations of reverse transcription PCR showing that 3R tau transcripts were absent in *MAPT*^S305N;Int10+3;S320F^ mice and minimally present in *MAPT*^P301S;Int10+3;S320F^ mice (**Figure 1F, G**). Likewise, a small amount of 3R tau transcripts was detected in *MAPT*^P301S;Int10+3;S320F^mice and was absent in *MAPT*^S305N;Int10+3;S320F^ mice in the real-time PCR analysis whereas the total tau expression levels were comparable among all groups (**Figure 1H**). We also observed similar pattens of 3R/4R tau ratios in the *MAPT*^S305N;Int10+3;S320F^ and *MAPT*^S305N;Int10+3;S320Y^ mice (**Supplementary Figure 2A-D**). These results indicate that the 10+3 G>A Intronic mutation affects the alternative splicing, causing a shift from 3R to 4R tau even in the presence of other mutations, and that the S305N mutation strengthens this effect in a manner described previously^5^.

### Spatiotemporal mapping of AT8-positive phosphorylated tau accumulation

To investigate the extent of phosphorylated tau distribution, we conducted immunofluorescence staining with AT8 antibody, which detects phosphorylation of tau at Ser202 and Thr205, on four different coronal sections (from anterior to posterior). The heat maps created by the intensity of AT8-positive signals revealed that the strongest tau pathology was observed in the hypothalamus and amygdala, with relatively less intensity in the somatomotor, somatosensory, auditory, retrosplenial and visual cortices in addition to the hippocampus in *MAPT*^P301S;Int10+3;S320F^mice at the age of 6 months (**Figure 2A-C**). We also observed intense AT8-positive signals in the piriform cortex, lateral septal nucleus, thalamus, entorhinal cortex, and midbrain areas. The *MAPT*^S305N;Int10+3;S320F^ mice showed a similar pattern of AT8-positive signals with a lower intensity (**Figure 2C**). The accumulation of phosphorylated tau increased progressively (and almost linearly) in an age-dependent manner both in the *MAPT*^P301S;Int10+3;S320F^ and *MAPT*^S305N;Int10+3;S320F^ mice (Figure 2D, E). Although it is not possible to determine the age of onset in pathological terms, extrapolation of the data shown in **Figure 2E** implies that tau accumulation is likely to start at 4-5 months of age or even earlier particularly in the hypothalamus. *MAPT*^S305N;Int10+3;S320Y^ mice also showed phosphorylated tau accumulation in the hypothalamus and amygdala in particular at the age of 6 months, but the extent of tau pathology was less than that of the *MAPT*^P301S;Int10+3;S320F^mice (**Supplementary Figure 2E,F**).

**Figure 2.**
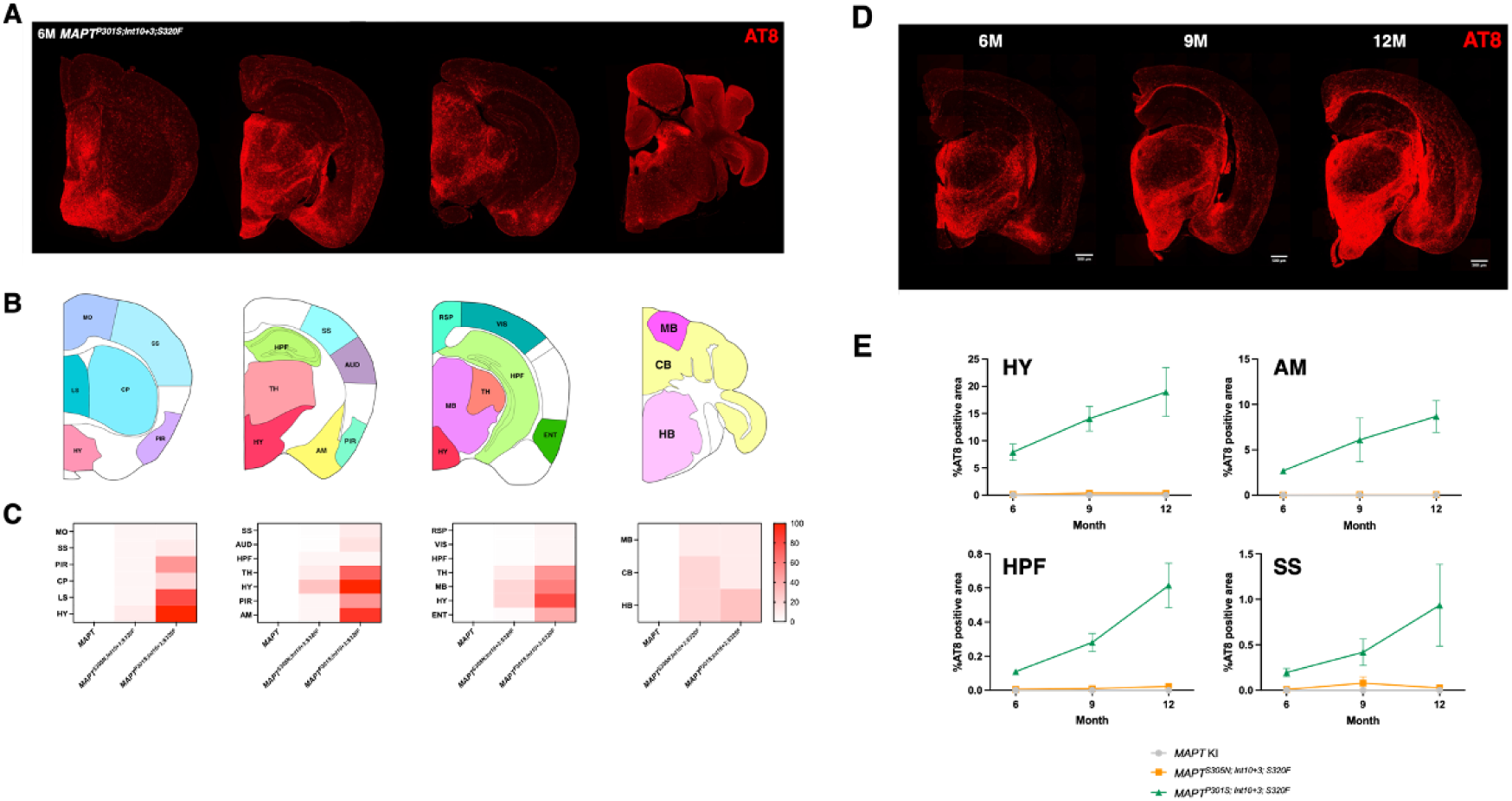
Spatio-temporal mappings of tau pathology in triple mutant mouse lines. **A.** Representative coronal sections of *MAPT^P301S;Int10+3;S320F^* mice stained with AT8 antibody. **B.** Classification of brain regions into several large functional areas. MO: somatomotor area, SS: somatosensory area, PIR: piriform area, CP: caudoputaman, LS: lateral septal nucleus, HY: hypothalamus, AUD: auditory area, HPF: hippocampal formation, TH: thalamus, AM: amygdala, RSP: retrosplenial area, VIS: visual area, MB: midbrain, ENT: entorhinal area, CB: cerebellum, HB: hindbrain. **C.** Heat maps of AT8 immunopositivity in different brain regions for *MAPT, MAPT^S305N;Int10+3;S320F^*, and *MAPT^P301S;Int10+3;S320F^* mice. **D.** Representative AT8 immunostained brain slice images of *MAPT^P301S;Int10+3;S320F^* mice at 6, 9 and 12 months of age. **E.** Time curves of AT8 immunopositivity for *MAPT, MAPT*^S305N;Int10+3;S320F^, and *MAPT*^P301S;Int10+3;S320F^ mice at HY, AM, HPF and SS. All immunostaining images were captured simultaneously with consistent contrast settings.

### Histological studies of tau pathology in triple mutant KI mice

To assess the pathological status of tau protein in the triple mutant *MAPT* KI mouse lines, we performed immunofluorescence analysis using a panel of antibodies that recognize the phosphorylated, oligomeric and pathological conformation structure of tau protein (**Figure 3A**). Immunohistochemistry using CP13 (pS202), PHF-1 (pS396/S404), TOC1, T22 and MC1 ^19–23^ antibodies clearly highlighted the presence of hyperphosphorylated and oligomeric/fibrillar tau in the brain of *MAPT*^P301S;Int10+3;S320F^ line (**Figure 3A, B**). With the identical panel of antibodies, we observed less intense and milder signals in *MAPT*^S305N;Int10+3;S320F^ and *MAPT*^S305N;Int10+3;S320Y^ mice (**Figure 3B** and **Supplementary Figure 3**). These results were consistent with those seen with DAB (3,3’Diaminobenzidine) staining (**Supplementary Figure 4**). The AT8 immunoreactivity colocalized with MAP2 and kinesin immunoreactivities (**Figure 3C**), indicating the presence of pathological tau in the neuronal dendrites, soma and axons.

**Figure 3.**
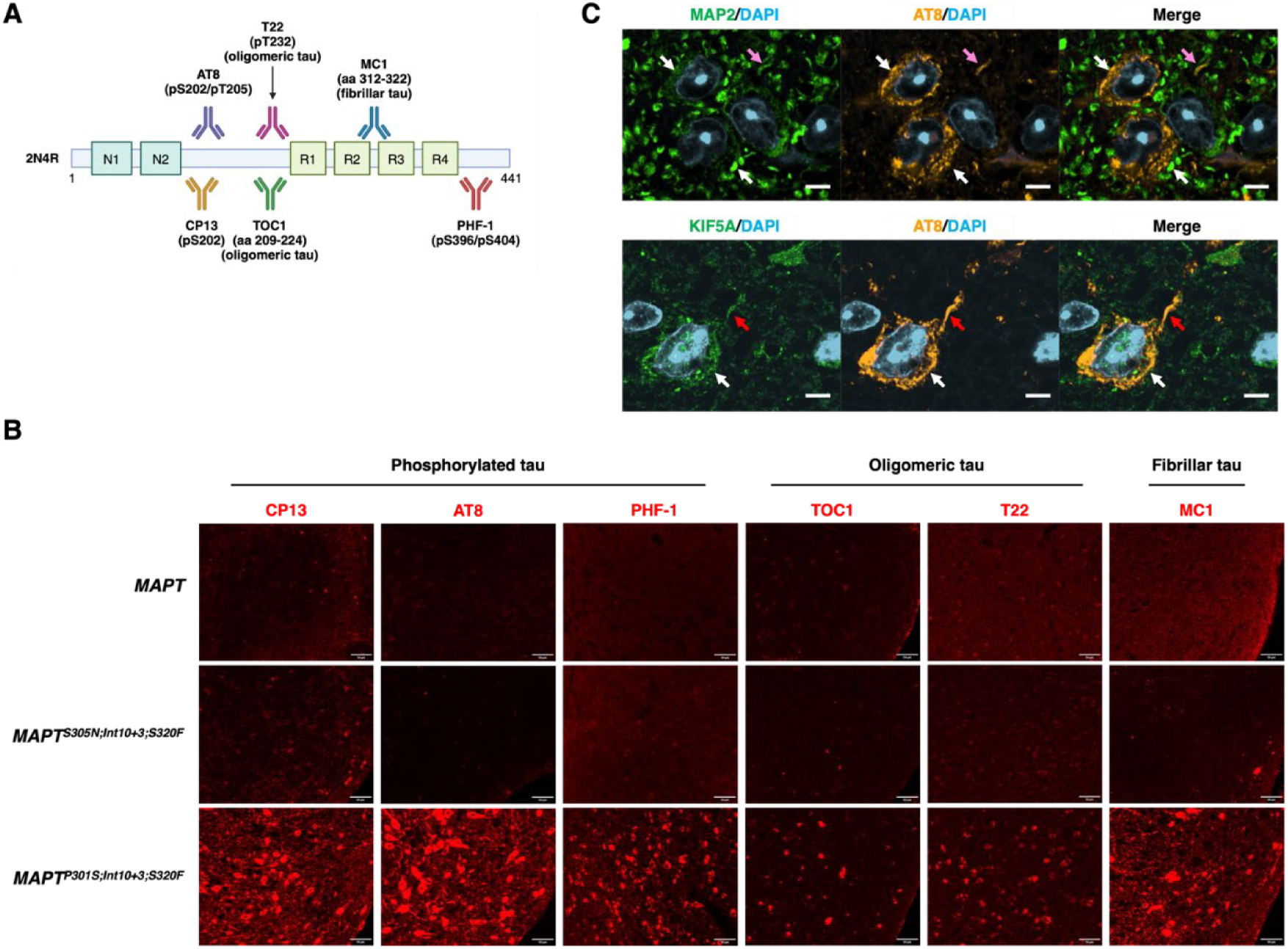
Immunohistochemical analysis of triple mutant *MAPT* KI mouse lines. **A.** Epitope maps of anti-tau antibodies used. **B.** Immunohistochemical analysis of hypothalamus of *MAPT, MAPT*^S305N;Int10+3;S320F^ and *MAPT*^P301S;Int10+3;S320F^ KI mice using CP13, AT8, PHF-1, TOC1, T22 and MC1 antibodies. Scale bars: 50 μm. **C.** Colocalization of AT8 signals with MAP2 (a marker for soma and dendrites; top panel) or KIF5A (a marker for soma and axons; bottom panel) in the cortical region of 12-month-old *MAPT*^P301S; Int10+3; S320F^ mice. Scale bars: 20 µm.

### Biochemical analysis of tau pathology in triple mutant KI mice

Sarkosyl fractionation is employed to isolate pathological tau proteins from bulk brain tissues ^24^. Through this method, we obtained Tris-buffered saline (TBS)-soluble fractions (S1), which contain physiological forms of tau, and sarkosyl-insoluble fractions (P3), which contain pathological forms of tau (i.e. tau protein with an oligomerization-prone conformation) (**Figure 4A**). In the S1 fractions, the expression level of total tau, represented by Tau13 antibody, showed distinct band patterns in the two mutant lines compared to wild-type human *MAPT* KI (**Figure 4B**). However, the densitometric analysis showed the amount of total tau was equivalent across genotypes (**Figure 4B and C**). Intriguingly, phosphorylation levels of tau, detected with AT8, CP13 and PHF-1 immunoreactivities, were significantly reduced in *MAPT*^S305N;Int10+3;S320F^ and *MAPT*^S305N;Int10+3;S320Y^ mice compared to the *MAPT* KI (**Figure 4B, C** and **Supplementary Figure 2G,H**). Sarkosyl-insoluble phosphorylated tau was detectable in both *MAPT*^P301S;Int10+3;S320F^ and *MAPT*^S305N;Int10+3;S320Y^ mice (**Figure 4D** and **Supplementary Figure 5**). To assess whether tau from triple *MAPT* KI mutants is capable of forming self-replicating assemblies (seeds), we employed biosensor cell lines overexpressing the repeat domain of tau containing the P301S mutation tagged with CFP or YFP (**Figure 4E**) ^25^. The brain lysates of S1 or P3 fractions from the hypothalamus and cortex in *MAPT*^P301S;Int10+3;S320F^ significantly increased seeding activity in these biosensor cells (**Figure 4F, G**), showing the presence of the pathological, seed-competent conformation of tau comparable to that of the PS19 line ^26^ even at the age of 6 months.

**Figure 4.**
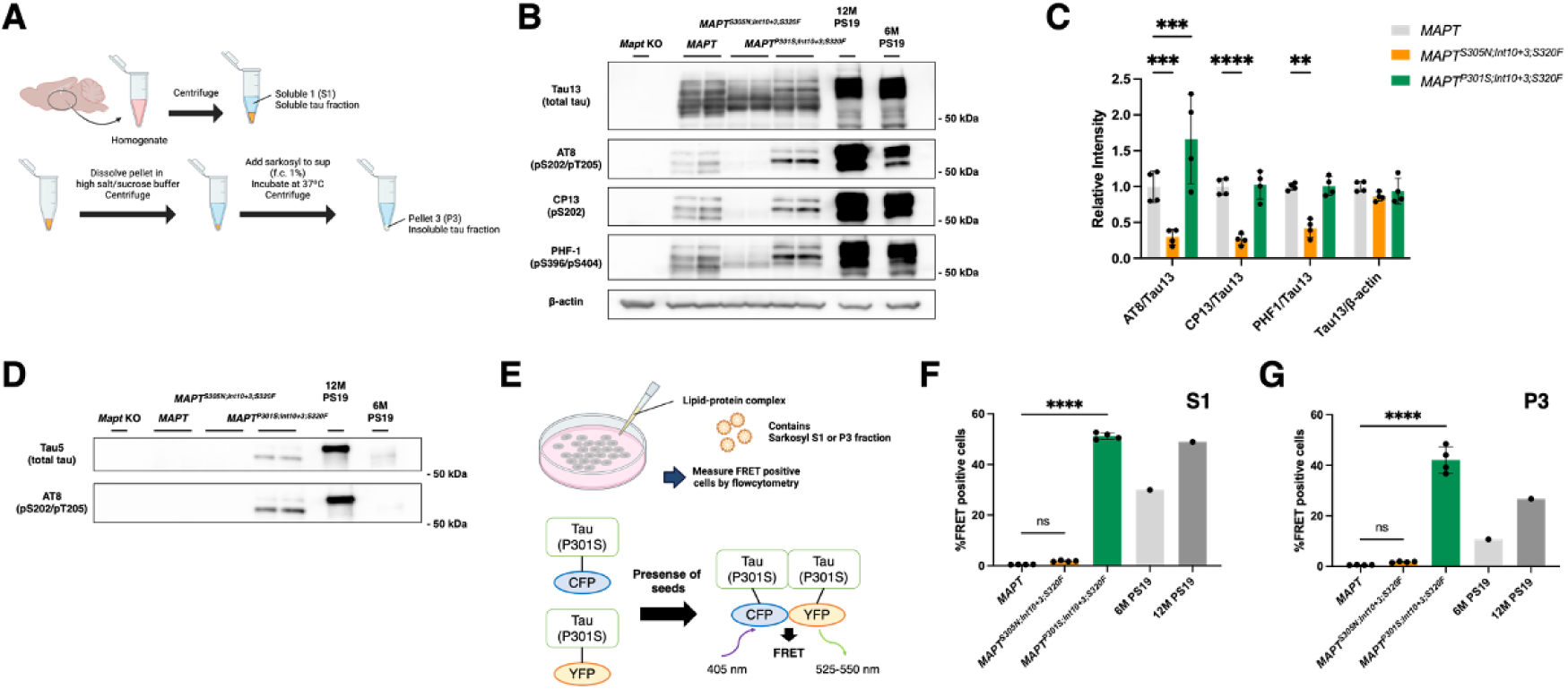
Sarkosyl fractionation of tau from brain extracts and their seeding potency analyzed by tau biosensor cells. **A.** A scheme for sarkosyl fractionation of tau. Sup: supernatant. **B, C.** Western blot analyses of TBS-soluble fractions from the thalamus of *MAPT*, *MAPT*^S305N;Int10+3;S320F^, *MAPT*^P301;Int10+3;S320F^ and *MAPT*^S305N;Int10+3;S320Y^ lines using Tau13, AT8, CP13 and PHF-1 antibodies (B) and densitometric quantification (C). *Mapt* KO mouse was used as a negative control. **D.** Western blot analysis of sarkosyl-insoluble fractions from the thalamus of *MAPT*, *MAPT*^S305N;Int10+3;S320F^ and *MAPT*^P301S;Int10+3;S320F^ lines using Tau5 and AT8 antibodies. **E.** Schemes for tau seeding assay using tau biosensor cells. **F, G.** The percentage of FRET-positive cells 48 h after introduction of S1 (F) or P3 (G) fractions to biosensor cells ^81^. Statistical significance was examined by two-way ANOVA (C). *: p<0.05, **: p<0.01, ***: p<0.005, ****: p<0.001.

### Neuroinflammation, synaptic loss and neurodegeneration of triple mutant *MAPT* KI mice

Neuroinflammation and synaptic loss are characterized as pathological hallmarks of AD and other tauopathies ^27^ ^28^. We thus performed immunohistochemistry of GFAP and Iba1 to assess neuroinflammation and found a slight upregulation of GFAP and no alterations in Iba1 immunoreactivity (**Figure 5A-C**). These results indicate that neuroinflammation is not proportionally exerted by tau accumulation, showing a reduced association of tau pathology with neuroinflammation in these models.

**Figure 5.**
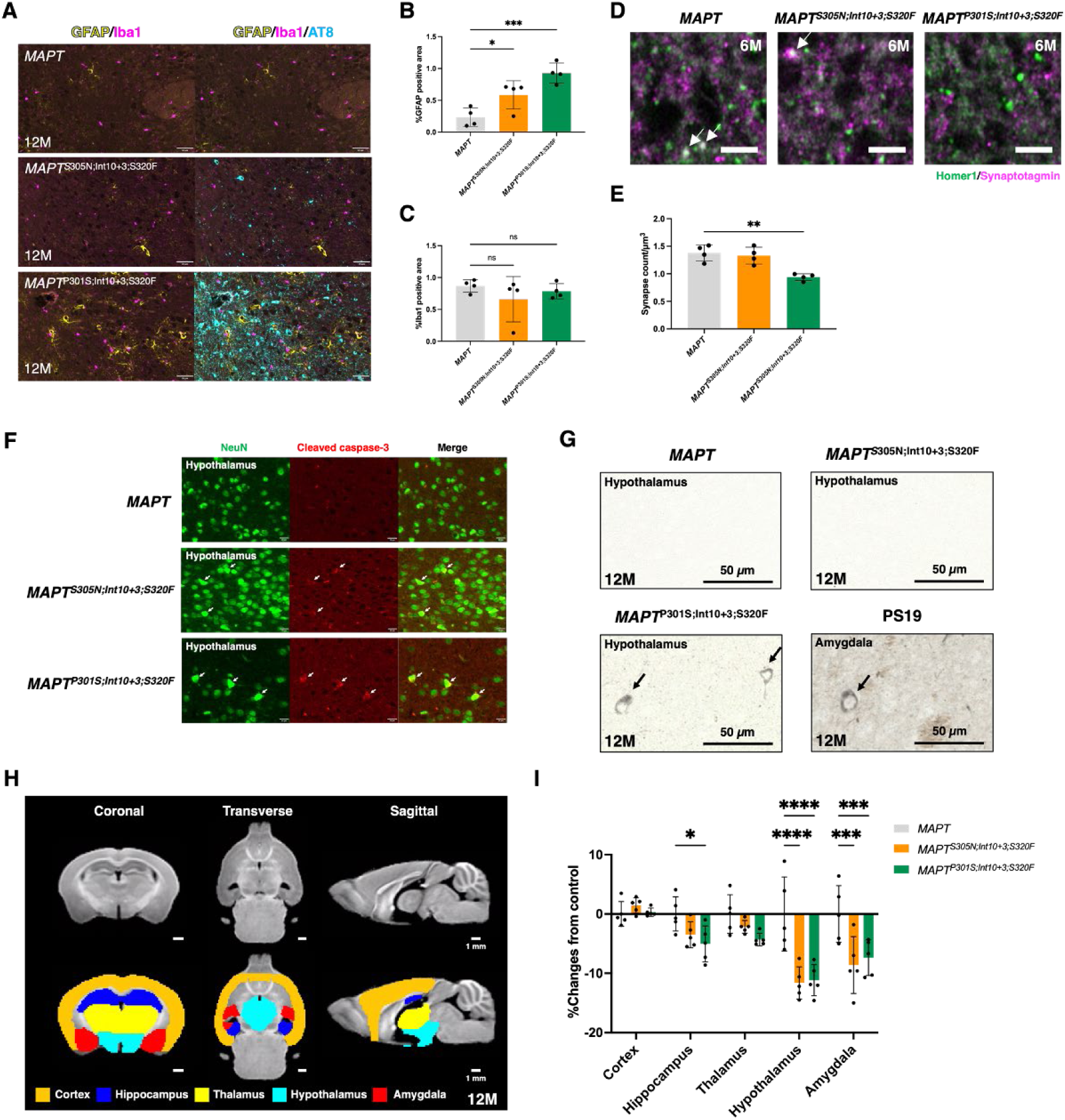
Neuroinflammation, synaptic loss and neurodegeneration in triple mutant *MAPT* KI mice. A-C. GFAP and Iba1 immunohistochemistry to assess neuroinflammation (A) and quantification of immuno-positive regions (B, C). **D, E.** Synapse colocalization analysis using a pre-synaptic marker (Homer1) and a post-synaptic marker (Synaptotagmin). **F.** Immunohistochemistry of an apoptosis marker cleaved caspase-3 with a neuronal marker NeuN. **G.** Gallyas staining-positive signals of *MAPT*^P301;Int10+3;S320F^ and PS19^26^. Scale bars: 50 µm. Statistical significance was examined by one-way ANOVA (B, C, E) or two-way ANOVA (G). *: p<0.05, **: p<0.01, ***: p<0.005. **H, I.** Regional analysis of T1 MRI images obtained from *MAPT, MAPT*^S305N;Int10+3;S320F^ and *MAPT*^P301;Int10+3;S320F^ lines (females, n=5 for each line).

Next, we analyzed the synaptic integrity using super-resolution confocal microscopy. Colocalization synaptic puncta detected by Synaptotagmin, a pre-synaptic marker, and Homer1, a post-synaptic marker, were significantly decreased in the hypothalamus of *MAPT*^P301S;Int10+3;S320F^ compared with those in *MAPT* mice (**Figure 5D, E**). No alterations of synaptic densities in the piriform cortices, whivh are relatively preserved from tau pathology, were seen in *MAPT*^P301S;Int10+3;S320F^ mice (**Supplementary Figure 6**). We also performed Western blot analysis of pre- and post-synaptic markers, synaptophysin and PSD-95. We found a trend toward decreased synaptophysin levels (p=0.08) and a significant increase in PSD-95 levels (p=0.0006) in the thalamic/hypothalamic fractions of brain extracts, whereas their levels in the cortical fractions remained unchanged (**Supplementary Figure 7**). These observations indicate an alteration of synaptic protein expression in the thalamus and hypothalamus.

Moreover, we observed immunopositivity of cleaved caspase-3 (cCasp3), an apoptosis marker, in hypothalamus (**Figure 5F**). NeuN, a neuronal marker, also colocalizes the cCasp3 signals at the age of 12 months; however, the NeuN signals were dense compared with cCasp3-negative cells, implying abnormal morphology of neurons. We then performed Gallyas staining, which highlights the presence of pathological tau accumulation, on these mice brains and found staining-positive cells in the hypothalami of 12-moths-old *MAPT*^P301S;Int10+3;S320F^ (**Figure 5G**).

We next performed magnetic resonance imaging (MRI) analysis to assess brain atrophy. Control images, which were obtained by averaging five *MAPT* control brain images, was divided into 20 sections, and five representative brain regions (cortex, hippocampus, thalamus, hypothalamus and amygdala) were chosen for comparison of brain volumes with mutant mice (**Figure 5H, Supplementary Figure 8**). Of those regions, the hypothalamus and amygdala brain volumes were significantly decreased in *MAPT*^S305N;Int10+3;S320F^ and *MAPT*^P301S;Int10+3;S320F^ mice in comparison with *MAPT* control mice, implying tau pathology-associated atrophy in the hypothalamus and amygdala (**Figure 5I, Supplementary Figure 8B**). In addition, a decrease in hippocampal volumes was also observed in *MAPT*^P301S;Int10+3;S320F^ mice.

In summary, GFAP is upregulated in the hypothalamus without changes to Iba1 immunopositivity, demonstrating the relevance of astrogliosis with abnormal tau. Accumulated gallyas-positive tau in the hypothalamus resulted in synaptic loss, apoptosis and atrophy, indicating a region-specific pathological tau-mediated neurodegeneration of *MAPT*^P301S;Int10+3;S320F^ mice in a manner consistent with the observations by MRI.

### Behavioral analyses in triple mutant KI lines with IntelliCage

The IntelliCage system is an automated apparatus that is to analyze a series of behaviors for group-housed mice ^29–32^. It contains four conditioning corners with two nosepoke holes each with a gate to access the drinking water spout (**Figure 6A**). This system automatically records access to the conditioning corners, nosepoking, and water licking of each mouse. Behaviors in three mouse lines, *MAPT* (control), *MAPT*^S305N;Int10+3;S320F^ and *MAPT*^P301S;Int10+3;S320F^ were analyzed. To assess whether the mutant lines have face validity as models of dementia especially with tauopathy, we investigated the signs of: (1) altered emotional responses, (2) repetitive/stereotypical behavior, (3) impairments in learning and behavioral flexibility, (4) impairments in behavioral inhibition and sustained attention, and (5) anhedonia and apathy.

**Figure 6.**
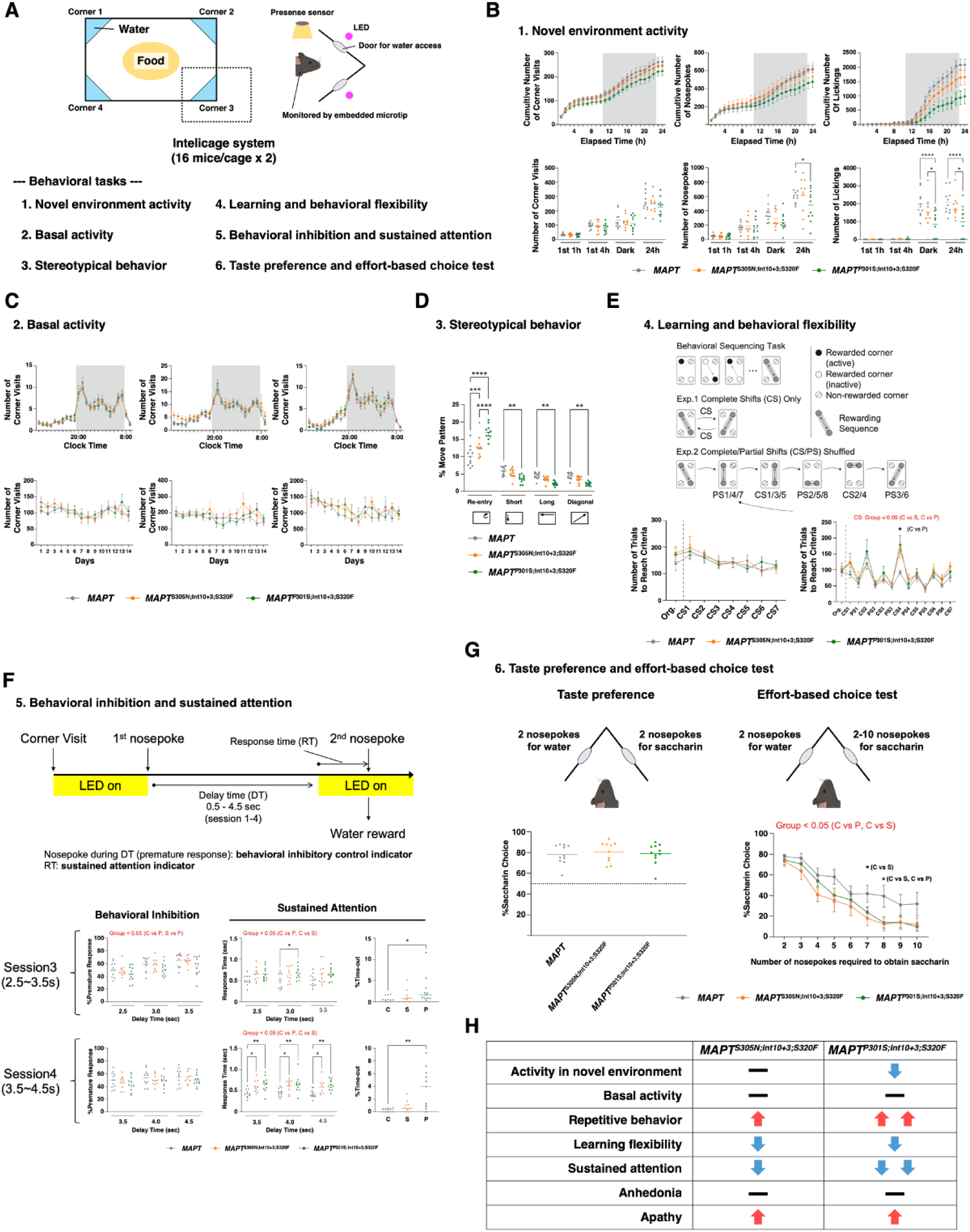
Behavioral abnormalities in triple mutant *MAPT* KI mouse lines observed using the IntelliCage system. **A.** Overview of the IntelliCage. Two IntelliCages were used to test a total of 32 female mice: *MAPT* (n=11), *MAPT*^S305N;Int10+3;S320F^ (n=10), *MAPT*^P301S;Int10+3;S320F^ (n=11), 5 or 6 mice per genotype per IntelliCage. **B.** Activity in a novel environment. Hourly cumulative number of corner visits, nosepokes, and lickings during the first 24 hours after the mice were introduced to the IntelliCages (upper panels) and those summaries (lower panels). **C.** Basal activity represented as daily averages over a 14-day period (upper panel) and as day-to-day total values (lower panels). **D.** Corner visiting patterns. Frequencies of (1) re-enteries to the same corner, corner-to-corner movements along (2) the shorter sides, (3) the longer sides, and (4) the diagonal. **E.** Learning and behavioral flexibility assessment. The upper panel presents the scheme of test protocol. Number of trials to reach criterion required for completing each phase of the CS-only session (lower left) and the CS/PS-shuffled session (lower right). **F.** Behavioral inhibition test and sustained attention test. Mice were required to withhold nosepokes during the delay time (DT). Nosepokes during the DT were recorded as premature responses. The response time (RT) between the second LED-on signal and the subsequent nosepoke was used as a measure of sustained attention. **G.** Sweet taste preference test and effort-based choice test. In both tests, one side of the nosepoke hole in each corner led to obtaining saccharin water, while the other side led to plain water. For the sweet taste preference test, the percentage of saccharin choice over two days are plotted. For the effort-based choice test, the percentage of saccharin choice are plotted against the number of nosepokes required to obtain saccharin water. **H.** Summary of behavioral phenotypes observed in mutant lines from the IntelliCage experiments. Statistical significance was examined by two-way ANOVA (B-G). *: p<0.05, **: p<0.01, ***: p<0.005, ****: p<0.001.

First, to assess the signs of altered emotional response of the mice, we analyzed their activity during the 24 hours after the mice were placed in the IntelliCage for the first time (**Figure 6B**). *MAPT*^P301S;Int10+3;S320F^ mice exhibited significantly decreased number of nosepokes and lickings compared to *MAPT* control mice. *MAPT*^S305N;Int10+3;S320F^ mice also exhibited a lower number of licking, while the effect was not significant (p=0.1836). These reductions were interpreted as a sign of elevated anxiety due to the exposure to a novel environment as the same indices in the following 14 days did not differ among groups (**Figure 6C**).

Second, we analyzed the signs of repetitive/stereotypical behavior. The corner preference and side preference of mutant mice were indistinguishable from controls (**Supplementary Figure 9A**). However, the percentage of re-entering the same corner was significantly increased in both *MAPT*^S305N;Int10+3;S320F^ and MAPT^P301S;Int10+3;S320F^ during the 14 days (**Figure 6D**). This indicated their sign of repetitive/stereotypical behavior, which is one of the clinical features of FTD^33^ (**Figure 6D**).

Third, to assess the ability of learning and behavioral flexibility, we conducted a task called SP-FLEX^34^. In this test, mice were required to learn a sequence of moving back and forth between two distant rewarding corners out of four corners and adapt to the subsequent serial rule shifts. For each individual, whenever the correct response rate reached a criterion (30%), the rewarding corners changed (i.e., rule shift). The number of trials to reach the criterion for the rule shift was used as the performance index. No genotype differences were observed in the “Complete Shifts (CS)-only” session, where two diagonal movement pattern shifts are repeated. However, in the more complicated “CS/Partial Shifts (PS)-shuffled” session, which is composed of six movement pattern shifts, both mutant lines required significantly more trials to reach the criterion throughout the session compared to control mice, indicating their difficulty in learning and behavioral flexibility (**Figure 6E**). We observed no changes in the number of nosepokes per corner visit in the two tests, showing normal activity of mice (**Supplementary Figure 4B**). See the **Methods** for detail experimental procedures.

Fourth, we assessed the ability of behavioral inhibition and sustained attention. In this test, mice had to withhold nosepokes for several seconds (delay time; DT), while sustaining attention on the LED lights, a signal for chance to acquire the water reward. A nosepoke during the DT are recorded as a premature response, and the reaction time (RT) to the LED lights after the DT serves as an indicator of the degree of sustained attention. A total of four sessions were conducted, with the DT extended as the sessions progressed. *MAPT*^P301S;Int10+3;S320F^ mice exhibited longer RT compared to *MAPT* control mice, indicating impairment in sustained attention. A delay in RT also emerged in *MAPT*^S305N;Int10+3;S320F^ mice when the DT was extended (session3 and 4) (**Figure 6F** and **Supplementary Figure 4C**).

Finally, to assess anhedonia and apathy-like behavior, we conducted the sweet taste preference test and the effort-based choice test using 0.1% saccharin water as hedonic reward. There were no genotype differences in the sweet taste preference test, indicating that both mutant lines had ability to perceive palatability (**Figure 6G, left**). On the other hand, in the effort-based choice test, the both mutant lines exhibited significantly lower preference for saccharin water compared to control when incremental efforts (i.e., the number of nosepokes) were required to obtain the reward (**Figure 6G, right**). This implied that the mutant lines had apathy-like traits, specifically as an elevated effort aversion.

In summary, *MAPT*^P301S;Int10+3;S320F^ mice exhibited signs of elevated anxiety, repetitive behavior, impaired learning and behavioral flexibility, impaired sustained attention, and apathy-like behavior, consistent with the clinical features of dementia with tauopathy (**Figure 6H**). A similar pattern of behavioral phenotypes was observed in the *MAPT*^S305N;Int10+3;S320F^ mice, though to a lesser extent. We also observed a body weight increase in *MAPT*^P301S;Int10+3;S320F^ mice, indicating increased food intake or abnormal metabolism (**Supplementary Figure 4D**).

### Effect of Aβ deposition on tau pathology in *MAPT* ^P301S;Int10+3;S320F^ mice

We crossbred *App*^NL-G-F^ mice with *MAPT*^P301S;Int10+3;S320F^ mice and obtained *App*^NL-G-F^ x *MAPT*^P301S;Int10+3;S320F^ mice to investigate whether *MAPT*^P301S;Int10+3;S320F^ mice are suitable for studying amyloid-β (Aβ) and tau interactions. Intriguingly, AT8 positivity was markedly enhanced by the existence of amyloid plaques, particulaly in the hippocampus and cortex (**Figure 7A,B**). However, the Aβ plaque density remained unchanged compared with *App*^NL-G-F^ (**Figure 7C,D**), indicating that tau pathology did not influence Aβ plaque formation. Furthermore, GFAP and Iba1 immunosignals were not different between *App*^NL-G-F^ and *App*^NL-G-F^ x *MAPT*^P301S;Int10+3;S320F^ (**Figure 7E-G**), suggesting that neuroinflammation was triggered by Aβ plaques regardless of the presence of tau pathology. Taken together, Aβ plaques, or their process of formation, direct abnormal tau to the hippocampus and cortex, and significantly accelerate tau pathology those regions.

**Figure 7.**
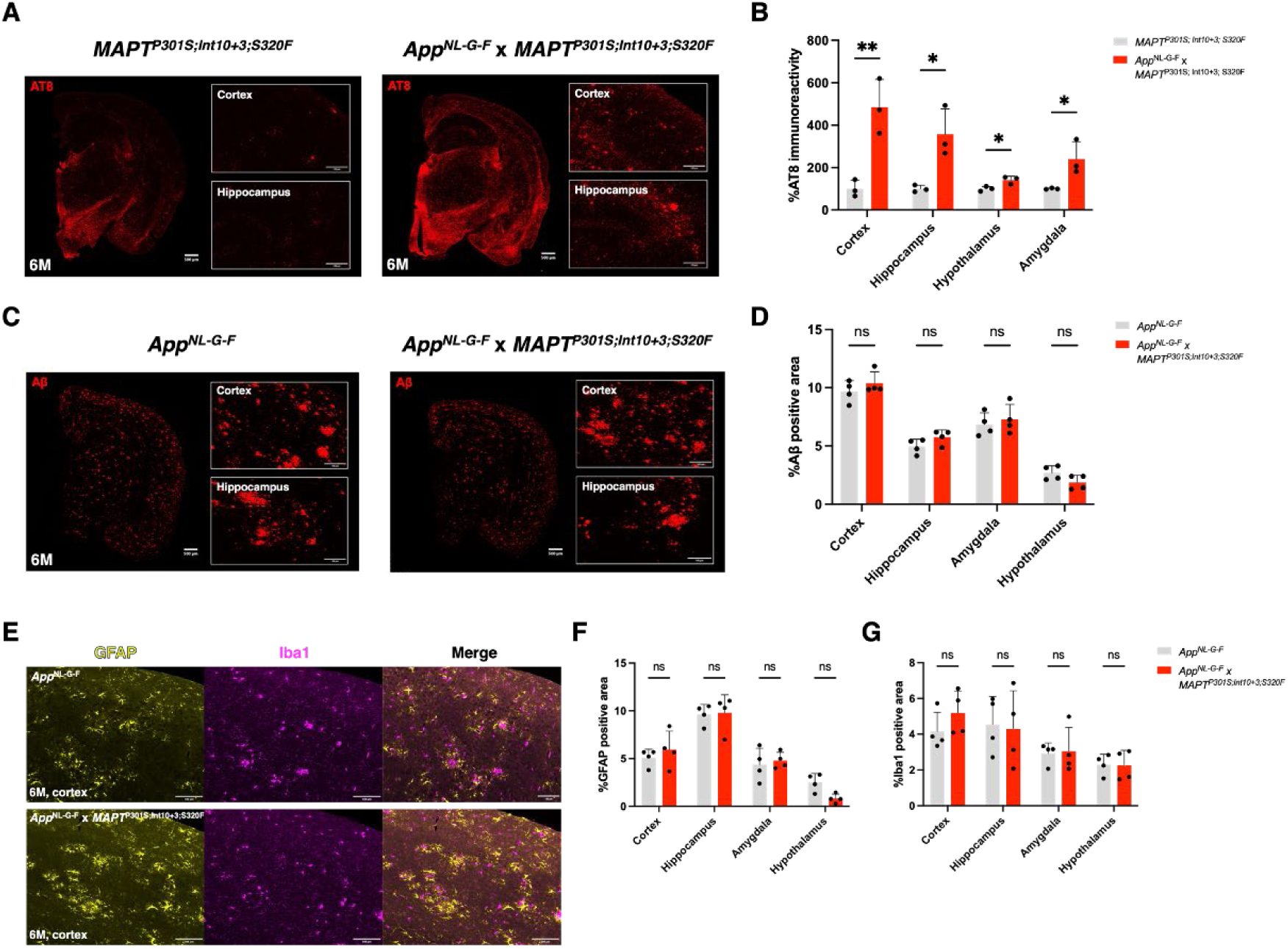
Effects of Aβ deposition on tau pathology in the hippocampus and cortex of *MAPT*^P301S;Int10+3;S320F^. **A, B.** Representative AT8 immunostaining images of brain sections from *MAPT*^P301S;Int10+3;S320F^ and *App*^NL-G-F^ x *MAPT*^P301S;Int10+3;S320F^ mice (A) and their quantification (B). **C, D.** Representative Aβ immunostaining images of brain sections from *App*^NL-G-F^ and *App*^NL-G-F^ x *MAPT*^P301S;Int10+3;S320F^ mice (C) and its quantification (D). **E-G.** Immunohistochemistry of neuroinflammation markers (GFAP and Iba1) in brain sections from *MAPT*^P301S;Int10+3;S320F^ and *App*^NL-G-F^ x *MAPT*^P301S;Int10+3;S320F^ mice (E). Regional quantification of GFAP (F) and Iba1 (G). Statistical significance was examined by one-way ANOVA (B, C, E) or two-way ANOVA (G). ns: not significant, *: p<0.05, **: p<0.01, ***: p<0.005.

## Discussion

*MAPT* mutations are causes of FTDP-17, but not AD. Nevertheless, the exclusively Aβ deposition-dependent tau pathology exhibited by *MAPT*^P301S;Int10+3^ mice crossbred with *App*^NL-G-F^ mice ^11^ demonstrates an evident cause-and-effect relationship between Aβ and tau pathologies. This is consistent with previous observations showing that massive tau pathology takes place in the cortex/hippocampus after Aβ deposition in aging human brains ^35–41^ and that transgenic APP or APP/PS1 overexpression accelerates tau pathology and neurodegeneration in mutant tau-transgenic mice ^42–48^. In this way, the mutant *MAPT* KI mice described in the present study should be highly suitable for preclinical studies of both FTDP-17 and AD. Because hippocampal tau pathology in *App*^NL-G-F^ mice crossbred with *MAPT*^P301S;Int10+3^ mice takes almost as long as 24 months ^11^, we decided to analyze tau pathology in *App*^NL-G-F^ mice crossbred with *MAPT*^P301S;Int10+3;S320F^ mice. The presence of Aβ deposition caused a significant increase of tau pathology in this mutant mouse line particularly in the hippocampus and cortex from as early as 6 months of age. While more detailed ananyses will be necessary, these bigenic mice will serve as a relevant model that reconstitutes the Aβ-tau axis observed in aging human brains.

In the present study, we have used a combination of multiple (triple) mutations in generating our FTD models, which does not happen in the real world, i.e. in clinical human cases. In this respect, our model mice are artificial, but we believe that this is one of the best approaches to reproducing the pathological phenotypes without inducing overexpression artifacts. In this way, we wanted to avoid the kind of overexpression artifacts that arose in the mutant APP and APP/PS1-overexpressing transgenic paradigms, most of which employed multiple mutations ^10,11^. (See Table 2 of Chapter 8 in Saido TC^11^.) These overexpression artifacts cause two uncertainties: [1] Uncertainty regarding why the artifacts arise and [2] Uncertainty regarding how the artifacts affect interpretation of the results obtained. Phenotypes such as endoplasmic stress, calpain activation and inflammasome involvement are likely experimental mirages. Indeed, use of the KI strategy resulted in no artificial phenotypes except for acceleration to the pathological state to our knowledge. However, choosing the right models becomes important in line with the purpose of animal experiments. For instance, presence of the Arctic mutation renders Aβ resistant to neprilysin-catalyzed proteolysis ^49^: *App*^NL-F^ or *App*^NL-F^ X *Psen1*^P117L^ lines are preferable to the *App*^NL-G-F^ line for studying Aβ metabolism^6,50^. Likewise, presence of the Swedish mutation perturbs β-secretase inhibitor action: the *App*^G-F^ line is preferable to the *App*^NL-G-F^ line for studying β-secretase inhibitors ^51^. We would like to emphasize here that the genome editing technology is applicable only mutant KI animals because transgene(s) are generally inserted into multiple loci in a random manner in the overexpressing transgenic animals ^52^.

In clear contrast to the *MAPT*^P301S;Int10+3;S320F^ mice, the *MAPT*^P301S;Int10+3^ mice that lack the S320F mutation exhibited minimal tau pathology up to 24 months of age ^11^, indicating that the combination of P301S and S320F mutations affects the oligomeric status of tau protein in a synergistic manner. It would be helpful to determine the 3D structure of tau protein in these animals by cryo-electron microscopy ^53–57^ as this may disclose how this particular combination of mutations affects the tau protein structure. Our findings are consistent with previous studies employing the combination of P301L/S and S320F mutations to show their synergistic effect on tau fibrillization using *in vitro*, transgenic or adeno-associated virus-dependent paradigms ^14–16^. We believe that one of the advantages of our KI models is the stably reproducible *in vivo* phenotypes.

The mutant *MAPT* KI mice that we have generated will be useful for preclinical *in vivo* studies of FTDP-17 as they are not affected by overexpression artifact-related phenotypes. Because tau protein expression in the new models is driven by the endogenous murine *MAPT* promoter, it may be possible to observe tau pathology in non-neuronal cells such as astrocytes ^58–61^ although more detailed pathological investigation will be necessary. Thus far, we have been unable to detect colocalization of pathological tau with GFAP.

The partial success of lecanemab and donanemab raised against Aβ oligomers and pyroglutamyl Aβ, respectively, in clinical trials to treat early AD ^62–65^ predicts that immunotherapies directed against pathological tau ^66–71^ may also be effective in treating FTDP-17, AD and other tauopathies, although none of the clinical trials have yet succeeded to our knowledge.

Finally, there will be more uses of these mutant *MAPT* KI lines in terms of cell biological applications. Establishment of embryonic stem (ES) cells or induced pluripotent stem (iPS) cells from the mutant mice will also make it possible to strategically generate drug candidates by, for instance, using the ES or iPS cell-derived neurons or organoids for *in vitro* drug screening. It would also be fascinating to apply Proteolysis-Targeting Chimera (PROTAC) technology to reduce the tau pathology load^72–76^.

## Materials and Methods

### Animals

All animal experiments were conducted in accordance with the guidelines of the RIKEN Center for Brain Science. C57BL/6J and ICR mice were used as zygote donors and foster mothers. C57BL/6J mice were also used for backcrossing with *MAPT* mutants. *MAPT* KI mice were generated as previously described (Hashimoto et al. 2019, Saito et al., 2019). *MAPT* ^P301S; Int10+3^ and *MAPT*^S305N; Int10+3^ KI mice were also obtained as described (Watamura et al., 2024, in press).

### Preparation of BE mRNA and sgRNAs

Base Editor (BE) which is a fusion protein composed of *Streptococcus pyrogenes* Cas9 (SpCas9) and rat APOBEC1 (apolipoprotein B mRNA-editing enzyme, catalytic polypepide-like1) (Komor et al., 2016) was used for the introduction of the FTD-associated mutation (S320F) into the *MAPT* gene of *MAPT* ^P301S; Int10+3^ and *MAPT*^S305N; Int10+3^ KI mice. For the synthesis of BE mRNA *in vitro*, plasmid vector pCMV-BE3 (Addgene plasmid #73021) was obtained from Addgene. To prepare the mRNA template for BE, PCR was performed with Herculase Ⅱ Fusion DNA Polymerase (Agilent Technologies #600675). BE was then synthesized with the mMESSAGE mMACHINE T7 Ultra Transcription kit (Thermo Fisher Scientific #AM1345). The template for the *in vitro* transcription of sgRNA was also synthesized by PCR with Herculase Ⅱ Fusion DNA Polymerase as previously described (Sasaguri et al.,2018). The MEGAshortscript T7 (Thermo Fisher Scientific #AM1354) and MEGAclear (Thermo Fisher Scientific #AM1908) kits were used for the *in vitro* transcription of sgRNA, while the CRISPR Design tool was used for creating sgRNA (Hsu et al., 2013). Oligonucleotide sequences used for the *in vitro* transcription of BE and sgRNA are described below.

BE: Forward 5’-GCCGCTAATACGACTCACTATAG-3’, Reverse 5’-GCCGCTAATACGACTCACTATAG-3’. sgRNA: Forward 5’-TAATACGACTCACTATAGGCTCCAAGTGTGGCTCATTGTTTTAGAGCTAGAA-3’, Reverse 5’-AAAAGCACCGACTCGGTGCCACTTTTTCAAGTTGATAACGGACTAGCCTTA TTTTAACTTGCTATTTCTAGCTCTAAAAC-3’.

### Microinjection of mouse zygotes

BE mRNA (60 ng/µl) and sgRNA (200 ng/µl) were injected into the cytoplasm of *MAPT* ^P301S; Int10+3^ and *MAPT*^S305N; Int10+3^ KI mice zygotes. After incubation at 37°C for 24 hours, embryos developed to the 2-cell stage were transplanted into host ICR mice.

### Off-target analysis

Off-target sites that accepted up to three mismatches were determined by COSMID (https://crispr.bme.gatech.edu/) ^18^. Target sites were amplified from tail genomic DNA by PCR using the Ex Taq-Polymerase kit (Takara #RR001A). Target sequencing was performed using a DNA sequencer (ABI 3730xl).

### Whole genome resequencing

#### Library construction

DNA samples extracted from mouse tail were prepared according to the Illumina TruSeq DNA sample preparation guide to obtain a final library of 300-400bp average insert size. 100 ng of genomic DNA were fragmented by the Covaris system with specific tubes, which generates double-strand DNA fragments with 3’ or 5’ overhangs. Fragmented DNAs were then converted into blunt ends using an End Repair Mix. The 3’ to 5’ exonuclease removes the 3’ overhangs, and the polymerase fills in the 5’ overhangs. Following the end repair, the appropriate library size was selected using different ratios of sample purification beads. Adenylation of 3’ ends was performed to prevent ligation during the adapter ligation reaction. A corresponding single “T” nucleotide on the 3’ end of the adapter provides a complementary overhang for ligating the adapter to the fragment. Multiple indexing adapters were ligated to the ends of the DNA fragments to prepare them for hybridizing onto a flow cell. PCR was used to amplify the enriched DNA library for sequencing. The PCR was performed with a PCR primer solution that annealed to the ends of each adapter.

#### Clustering and sequencing

For cluster generation, the library was loaded into a flow cell where fragments were captured on a lawn of surface-bound oligonucleotides complementary to the library adapters. Each fragment was then amplified into distinct clonal clusters through bridge amplification. When cluster generation was complete, the templates were ready for sequencing. Illumina SBS technology utilizes a proprietary reversible terminator-based method that detects single bases as they are incorporated into DNA template strands. As all four reversible, terminator-bound dNTPs were present during each sequencing cycle, natural competition minimized incorporation bias and greatly reduced raw error rates compared to other technologies. The result was a highly accurate, base-by-base sequencing that virtually eliminated sequence-context-specific errors, even within repetitive sequence regions and homopolymers.

#### Generation of raw data

The Illumina platform generates raw images and base calling through an integrated primary analysis software called Real-Time Analysis (RTA). The BCL (base calls) binary files were converted into FASTQ files using the Illumina package bcl2fastq2-v2.20.0.

#### Sequencing analysis

Resulting reads were aligned using Isaac Genome Alignment software (version 01.15.02.08), which is designed to align next-generation sequencing data with low-error rates ^77^, and Isaac Variant Caller (IVC) (version 2.0.13), which is used for identifying single-nucleotide variants (SNVs) and small insertions and deletions in the diploid genome case. The annotation tool SnpEff was used to categorize the effect of variants on genes ^78^.

#### Brain samples preparation

Mice were perfused with PBS for 2 min after anaesthesia. The brain was collected and sectioned along the midline to separate the two hemispheres. One side was immediately cut into five samples (cortex 1, 2, 3 from anterior to posterior, hippocampus, thalamus), frozen in liquid nitrogen, and stored at -80°C. The other side was incubated in 4% paraformaldehyde (Nacalai Tesque #09154-85) overnight at 4°C, washed in PBS, and embedded in paraffin for sectioning.

#### Sarkosyl fractionation

Sarkosyl fractionation was performed as described previously ^24^. Briefly, frozen brain tissues were weighed and homogenized in homogenization buffer (TBS supplemented with protease inhibitor and phosphatase inhibitor cocktail) with 0.1 g tissue weight/ml concentration, followed by ultracentrifugation (26,300 x g, 4°C, 20 min). The supernatant was used as the S1 fraction (TBS-soluble fraction). The pellet was dissolved in high salt/sucrose buffer (10 mM Tris-HCl pH 7.5, 0.8 M NaCl, 10% sucrose, 1 mM EGTA, protease inhibitor, phosphatase inhibitor) with a half volume of homogenization buffer, followed by ultracentrifugation (26,300 x g, 4°C, 20 min). Sarkosyl solution was added to the obtained supernatant for a 1% concentration, and the solution was incubated at 37°C for 1 hour. Finally, the P3 fraction (sarkosyl-insoluble fraction) was obtained by ultracentrifugation (150,000 x g, 4°C, 1 hour). P3 was dissolved in 0.5 µl/mg brain tissue Tris-EDTA buffer (pH 8.0) (TE buffer) (06890-54, Nacalai tesque) for Western blotting.

#### Western blotting

S1 and P3 fractions were subjected to sodium dodecyl sulfate-polyacrylamide gel electrophoresis (SDS-PAGE), followed by transfer to a PVDF membrane. The protein concentration of S1 fraction was measured by BCA assay to adjust the concentration. For detection of the tau isoform pattern, brain lysates were dephosphorylated with lambda-phosphatase (Santa Cruz Biotechnology #sc-200312A) according to the manufacturer’s protocol. Membranes were then treated with ECL prime blocking buffer (GE healthcare RPN418) for 1 hour and incubated with antibody at 4°C. Dilution ratios of antibodies are listed in **Supplementary Table 2**. Immunoreactive bands were visualized with ECL Select detection reagent (GE Healthcare RPN2235) and a LAS-3000 Mini Lumino image analyzer (Fujifilm).

#### Immunohistochemistry

Paraffin-embedded mouse brain sections were deparaffinized and applied to antigen retrieval by autoclaving sections at 121°C for 5 min in 0.01 M citrate buffer (pH 6.0). Sections were then rinsed in tap water, treated with 0.3% H_2_O_2_ in methanol to inactivate endogenous peroxidases, rinsed in water and TNT buffer (0.1M Tris-HCl pH 7.5, 0.15M NaCl, 0.05% Tween20), and blocked in blocking solution composed of 0.5% blocking reagent powder (AKOYA Biosciences, SKU FP1012) in TNB buffer (0.1M Tris-HCl pH 7.5, 0.15M NaCl). The sections were incubated overnight at 4°C with primary antibodies (Listed in **Supplementary Table 2**) diluted by TNB buffer and washed three times in TNT for 5 min, followed by incubation in biotinylated secondary antibodies diluted by TNB buffer at room temperature for 1 hour. After washing three times for 5 min, the sections were incubated in horseradish peroxidase (HRP)-conjugated streptavidin for 30 min, washed in TNT, and applied to tyramide-fluorescein isothiocyanate or tyramide-rhodamine solution for 10 min. Finally, the sections were stained with DAPI (Cell Signaling Technology #4083S) diluted in TNB buffer before mounting with ProLong Gold Antifade Mountant (Invitrogen, P36934). The mounted sections were scanned with a VS200 digital slide scanner (Evident).

#### MRI

MR images were acquired using a 7.0-T BioSpec 70/40 system with a ^1^H receive-only 4-channel phased array MRI CryoProbe (Bruker, Billerica, MA, USA) with the software for operation (ParaVision 360 Ver. 3.4; Bruker). T1-weigheted anatomical brain images were obtained 3D Modified Driven Equilibrium Fourier Transform (MDEFT) sequence with the following parameters; repetition time 3000 ms, echo repetition time = 10 ms, echo time = 2.0 ms, inversion delay = 1100 ms, field of view = 18 × 18 × 18 mm^3^, matrix size = 144 × 144 × 80 mm^3^, voxel size = 0.125 × 0.125 × 0.125 mm^3^, fat suppression = on, and number of average = 1. The scanning time was approximately 9 minutes for each mouse under inhalation anesthesia with 1.5% isoflurane. The acquired data were reconstructed and exported to the images using ParaVision software in DICOM format. Images were obtained by manually skull-stripping using PMOD software (ver. 4.1, PMOD Technologies LLC, Fällanden, Switzerland) to create the brain mask. The regional brain volume was measured utilizing a software package, the Advanced Normalization Tools following a published method^79^ The anatomical image was non-linearly registered to the mouse MR atlas R atlas^80^.

### Behavior analyses using the IntelliCage

#### Housing and testing environment

All behavior tests using IntelliCage (TSE Systems GmbH) were conducted at the animal facility of Phenovance LLC (Chiba, Japan) in accordance with that company’s Animal Care and Use Committee guidelines (Experiment Approval Number: PRT202405_52RS). The animal facility was maintained at 20-26°C, 40-70% relative humidity, 10-25 ventilation per hour, and a 12 h light-dark cycle (8:00 ON, 20:00 OFF). Two weeks before starting the IntelliCage experiments, mice were anesthetized by isoflurane inhalation with the aid of an SN-487-1T Scan instrument (SHINANO manufacturing) and implanted with a small, glass-covered radio-frequency identification (RFID) tag via a disposable injector with needle (2.12 x 12 mm, FDX-B ISO11784/11785, FAREAD Technology). The mice were housed in conventional plastic cages (2000P type, 612 x 435 x 216 mm, Tecniplast) in groups of 12-13 animals until the experiment.

#### IntelliCage System

IntelliCage is a fully automated testing apparatus for monitoring spontaneous and cognitive behaviors of group-housed RFID-tagged mice in a home cage (Kiryk et al. 2020). The corners of the cage are equipped with four triangular operant chambers (hereafter referred as corner A, B, C, and D) that permit the entry of a single mouse at a time. The RFID antenna at each corner entrance recognizes the tag ID implanted in each mouse, while infrared sensors and licking sensors inside the corner chambers monitor mouse behavior. Each corner holds two water bottles, the nozzles of which can be accessed through two nosepoke holes with motorized doors; the infrared beam-break response detector on the nosepoke hole responds to the mouse nosepokes which trigger door opening/closing. A small concrete block was placed in front of each corner as a footstool to make it easier for mice to enter the corner. Experimenters can flexibly program the rules for opening/closing of the doors and the lighting pattern of three LED lights located above each nosepoke hole. Two IntelliCages were used for experiments and each group of mice was divided as evenly as possible (5-6 mice per genotype per IntelliCage) for a total of 16 mice per IntelliCage. Each IntelliCage was usually cleaned once a week, at which time the general health of each mouse was also carefully checked (body weights were measured; condition of skin, hair, eyes, mouth, tail and basic walking movements in an open cage were visually assessed. During the experiment, each IntelliCage was placed in a soundproof wooden isolation box (Phenovance LLC) for environmental control.

#### Programs used for IntelliCage experiments

For the following IntelliCage experiments, each task program was selected from the Phenovance Task Library, which contains a collection of task programs for IntelliCage created by Phenovance LLC using specialized software (Designer software, TSE Systems).

#### Exploratory behavior, basal activity, repetitive/stereotypical behaviors

On the first day of the IntelliCage experiment, mice were transferred from the conventional 2000P cages to IntelliCages by “tunnel handling” (Hurst and West 2010), two hours into the light period (10:00 AM). All doors in the corner chambers were kept open and mice were free to drink water from any corner. The number of corner visits, nosepokes, and lickings during the day were measured as indices of exploratory behaviors in novel environment. In the following 14 days, all the doors were closed and a single nosepoke opened the door for 3000 ms to allow mice to drink water. The number of corner visits, nosepokes, and lickings from the 14 days were used to calculate the indices of basal activity level and repeptitive/stereotypical behaviors (corner visiting patterns, corner preferences, side preferences).

#### Self-paced Learning and Behavioral Flexibility Test (SP-FLEX)

This test is designed to assess the learning ability of mice to the behavioral sequence rule and behavioral flexibility to adapt to repetitive rule shifts (modified from an original protocol developed in a previous study by Endo et al., 2011). Each mouse was assigned a pair of corners as rewarding corners and had to shuttle between them to obtain water as a reward; water could not be obtained from the other two corners (non-rewarding corners) until the rule switched. When a mouse nosepoked at one of the rewarding corners, the door opened for 2000 ms to permit the water intake, with the following restrictions: 1) The door opened only once per visit, and 2) a mouse was not permitted to obtain the reward by consecutively re-entering the same corner. Because of these restrictions, mice were forced to go back and forth between the assigned two rewarding corners in order to obtain the reward, one defined as the active corner and the other as the inactive corner. A success response was defined as obtaining a reward by a direct move from the previously active rewarding corner to the current active rewarding corner, while a failure response was defined as visiting one of the two non-rewarding corners and applying at least one nosepoke. The success and failure responses were automatically detected and input to Wald’s sequential probability ratio test (SPRT) statistics to calculate each mouse’s probability of success. The significance level for the acceptance of each criterion was 0.05. Corner visits without nosepokes were omitted from the SPRT calculation. Whenever the performance of a mouse reached the pre-set upper criterion (30%), the assignment of rewarding and non-rewarding corners for that mouse switched. Note that the switch only occurred for mice whose success rate reached the upper criterion of SPRT. The test consisted of two sessions called the “CS (complete shift)-only” session and the “CS (complete shift) / PS (partial shift)-shuffled” session, each of which lasted 24 hours per day for ten days. The detailed protocol for each session is described below (see also a previous study conducted using the same protocol) (Balan et al. 2021).

Session 1. CS (complete shift)-only session: In this session, the mice were required to learn a behavioral sequence in which they were rewarded for moving back and forth between two diagonally opposite corners (e.g., corners A, C) out of four corners (corners A, B, C, D). When the performance reached the upper criterion of SPRT, the two corners that had been rewarding (corners A and C) and the two non-rewarding corners (corners B and D) were reversed. Note that the term “complete shift” or “CS” means that all the previously rewarding corners and all the previously non-rewarding corners were reversed at the same time. Within a session, there were multiple test phases: The Original learning phase (Org) is the phase of initial learning of the behavioral sequence rule until the first reversal occurs; Org was followed by the CS1 phase, when the rewarding and non-rewarding corner assignments reversed for the first time; the phases proceed every time the reversal occurs (e.g., CS2, CS3, and so on).

Session 2. CS (complete shift) / PS (partial shift)-shuffled session: In this session, the same protocol as in the CS-only session was used until the end of the CS1 phase. When a mouse reached the upper criterion of SPRT in the CS1 phase, only one of the two rewarding corners was replaced by one of the two non-rewarding corners (henceforth referred to as “partial shift” or “PS”). After that, the CS and PS phases were alternately imposed on each mouse. All combinations of the two corners out of four corners were presented twice each.

The number of trials, which is defined as the sum of the success and failure responses required to reach the upper criterion, was used to assess the efficiency of learning and behavioral flexibility at each task phase. In addition, the average numbers of nosepokes per active rewarding corner, inactive rewarding corner, and non-rewarding corners were calculated as secondary indices of behavioral flexibility. This is based on the notion that no matter what type of corner visit it is, more than a single nosepoke is redundant in this task: The second nosepoke will not provide any further reward in the active rewarding corner, and neither the inactive rewarding corner nor the non-rewarding corner provide reward, regardless of how many nosepoke attempts a mouse may make. Therefore, repetitive nosepokes in each corner visit can be interpreted as an inability to flexibly terminate the meaningless actions; in other words, behavioral inflexibility derived from impulsive or perseverative characteristics.

#### Behavioral Inhibition and Sustained Attention Test

This test was designed based on the protocol of an operant schedule called differential reinforcement of low rates with limited hold (DRL-LH): the protocol assesses whether mice could maintain impulsive nosepoking behavior under control for a certain amount of time and whether they could sustain attention during that time. In this test, all mice were permitted to obtain water as a reward in any of the four corners by making nosepokes in a way that followed the rules below: When a mouse entered a corner, the leftmost and rightmost LED lights above nosepoke holes were lit yellow. The LED lights turned off when the mouse made its first nosepoke on either side, but the door did not open. Then, after a certain amount of time (delay time; DT), the LED lights turned back on (reward signal), at which time the mouse could obtain a reward by making its second nosepoke. The door opened for 4000 ms in response to the nosepoke; after it closed, the trial for that corner visit ended, meaning that the mouse had to leave the corner and re-enter it for more reward.

On the first day of the training phase (Training Day 1), the delay time was set to 0, so mice were rewarded for nosepoking twice in a row. On Training Day 2, the delay time was set to 100 ms, on Training Day 3 to 200 ms, and on Training Days 4-5 to 500 ms. If the mouse nosepoked during the delay time, the timer was set back to zero, meaning that mice had to suppress nosepoke activity during the delay time in order to obtain rewards in an efficient manner.

At the end of Training Day 5, the following four test sessions were conducted (four days per session): In Session 1, a delay time of 500 ms, 1000 ms, or 1500 ms was presented randomly in each trial. If a nosepoke was made during the delay time, the timer was set back to zero and recorded as one premature response. In the analysis, the percent of premature responses during each delay time was used as an indicator of the difficulty of behavioral inhibitory control. A nosepoke during the reward signal opened the door for 4000 ms to permit mice to drink water as a reward. However, the reward signals were presented only for five seconds, and if no nosepoke was made during that time, the reward opportunity was lost for that trial. The response time between the presentation of the reward signal and the nosepoke was used as a measure of sustained attention. In Session 2, random DTs of 1000 ms, 1500 ms, or 2000 ms were applied. In Session 3, random delays of 1500 ms, 2000 ms, or 2500 ms were applied. In Session 4, random delays of 2500 ms, 3000 ms, or 3500 ms were applied. Otherwise, the rules were the same as in Session 1. For each of Sessions 1, 2, 3 and 4, data from the second day onward were used to evaluate performance, as the first day of each four-day session required familiarization with the new delay time condition.

#### Sweet Taste Preference and Effort-based Choice Test

0.1% saccharin (199-08665, FUJIFILM Wako Pure Chemical Corporation) water was used as the hedonic reward, preferred by the C57BL/6 strain.

In the sweet taste preference test and the subsequent effort-based choice test, mice were able to freely drink saccharin or plain water from any of the four corners; in two corners, they were able to drink saccharin water from the right nosepoke holes and plain water from the left. In the remaining two corners, the positions of the left and right bottles were reversed.

The protocol for the sweet taste preference test was as follows. When a mouse visited a corner, the doors of the left and right nosepoke holes were automatically opened. When the mouse nosepoked on either side, the door on the side of the nosepoke closed after 1500 ms (henceforth called the “tasting trial”). After the tasting trial, when the mouse nosepoked the closed door again, the door opened for 2000 ms (hereafter called “preference-based choice”). The mouse could repeat the preference-based choice as many times as it wanted in a single corner visit. In the analysis, the percentage of preference-based choices made for saccharin water out of the total choices made for both types of solutions was used as a sweet taste preference index. The test was conducted over two days. Day 1 began at 12:00 noon, four hours after the start of the light period, with the saccharin and plain water bottles inserted into the IntelliCage. On the second day of the test, the positions of the bottles on the left and right sides of each corner were reversed at 12:00 noon. The mean value of the preference-based choice rate for the saccharin water on each day was used as a representative value for comparison between groups. After confirming that no significant difference in preference for the 0.1% saccharin water was apparent between the groups, the degree of motivation for acquiring this sweet reward was tested using the PR operant schedule. In this test, as in the previous sensitivity test protocol, the mice were able to choose between the saccharin solution and plain water at each corner, but the following rules were added: After the tasting trial, mice were intially required to make two consecutive nosepokes to receive a 4000 ms drink of either saccharin water or plain water, at the beginning of the effort-based choice task. Thereafter, each time the total number of preference-based choices exceeded 50, the number of nosepokes required to drink saccharin water was increased one by one until 10 nosepokes were required for mice to obtain an opportunity to drink saccharin water. During this period, the rates of preference-based choice for saccharin water to total preference-based choice was used in the analysis as an index of motivation to seek palatable pleasure.

## Acknowledgements

We thank Ryo Fujioka and Naoko Mihira at RIKEN CBS, Shoko Uchida and Akira Sumiyoshima at QST for technical advice. We also express our gratitude to Yukiko Nagai-Watanabe for her secretarial assistance. This work was supported in part by research grants from the RIKEN Aging Project (T.C.S.) the Strategic Research Program for Brain Sciences and the Japan Agency for Medical Research and Development (AMED) under JP20dm0207001 (T.C.S.), JSPS KAKENHI Grant Number 21K15378 (N.W.) and Overseas Research Fellowships (N.W.), and the RIKEN FY2020 Incentive Research Project (N.W.).

## Data and material availability

The datasets generated and/or analyzed during the current study are available from the corresponding authors upon reasonable request. Manuscripts of publications in press cited in this paper are also available upon request. The mutant mice (listed in **Table 1** and **Supplementary Table 2**) will be made available to non-profit organizations for non-profit research free of charge (except for shipping and handling costs) upon request to TCS. We ask for-profit organization(s) to contact TCS on this matter.

## Author contributions

NW, MQ and NM generated and bred all the mutant *MAPT* KI mice. TM, MQ, NK, HS, SB, MF, KD, SB, TE, HH, HK, AM, HM, NS, MS, MH and NW planned, performed, analyzed experiments: they also discussed the results thoroughly. TM, MQ, NK, SB, TE, HM, MS and NW created the figures. TM, MQ, NW and TCS wrote the manuscript with the assistance of the other authors.

Saido 241114 related manuscript

A paper in press for publication in Nature Neuroscience, which was too heavy to upload to your link, can be downloaded from the following link.

https://www.dropbox.com/scl/fi/wf2kktpvrak5k4vt6f6qu/Saido-241014-Manuscript-light.pdf?rlkey=5pbxwiciu88v64ykevw70e8w1&st=854e1n9n&dl=0

## Supplementary Information

### Supplementary Figures

**Figure S1.**
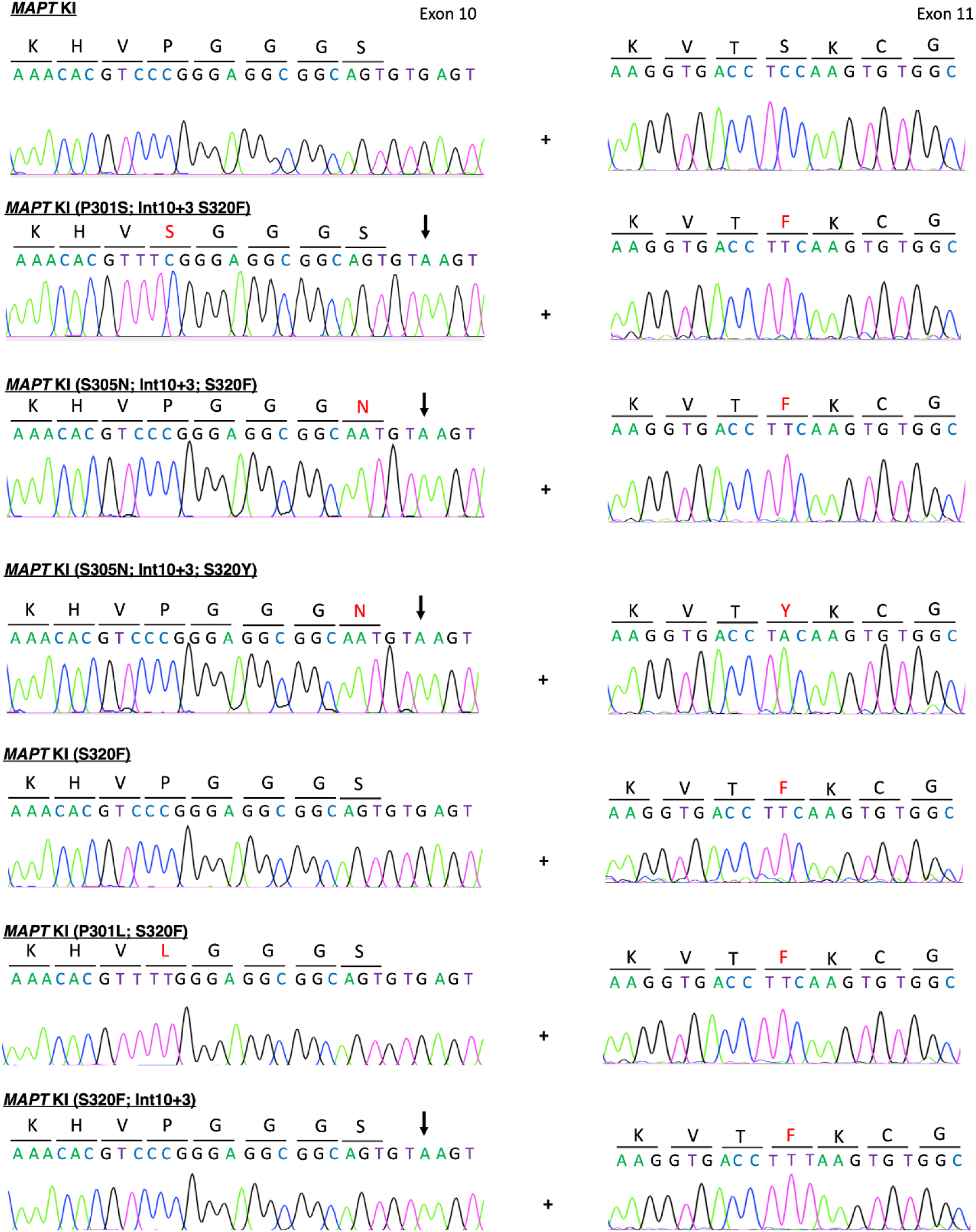
Sanger sequence data for the panel of mutant *MAPT* KI mouse lines.

**Figure S2.**
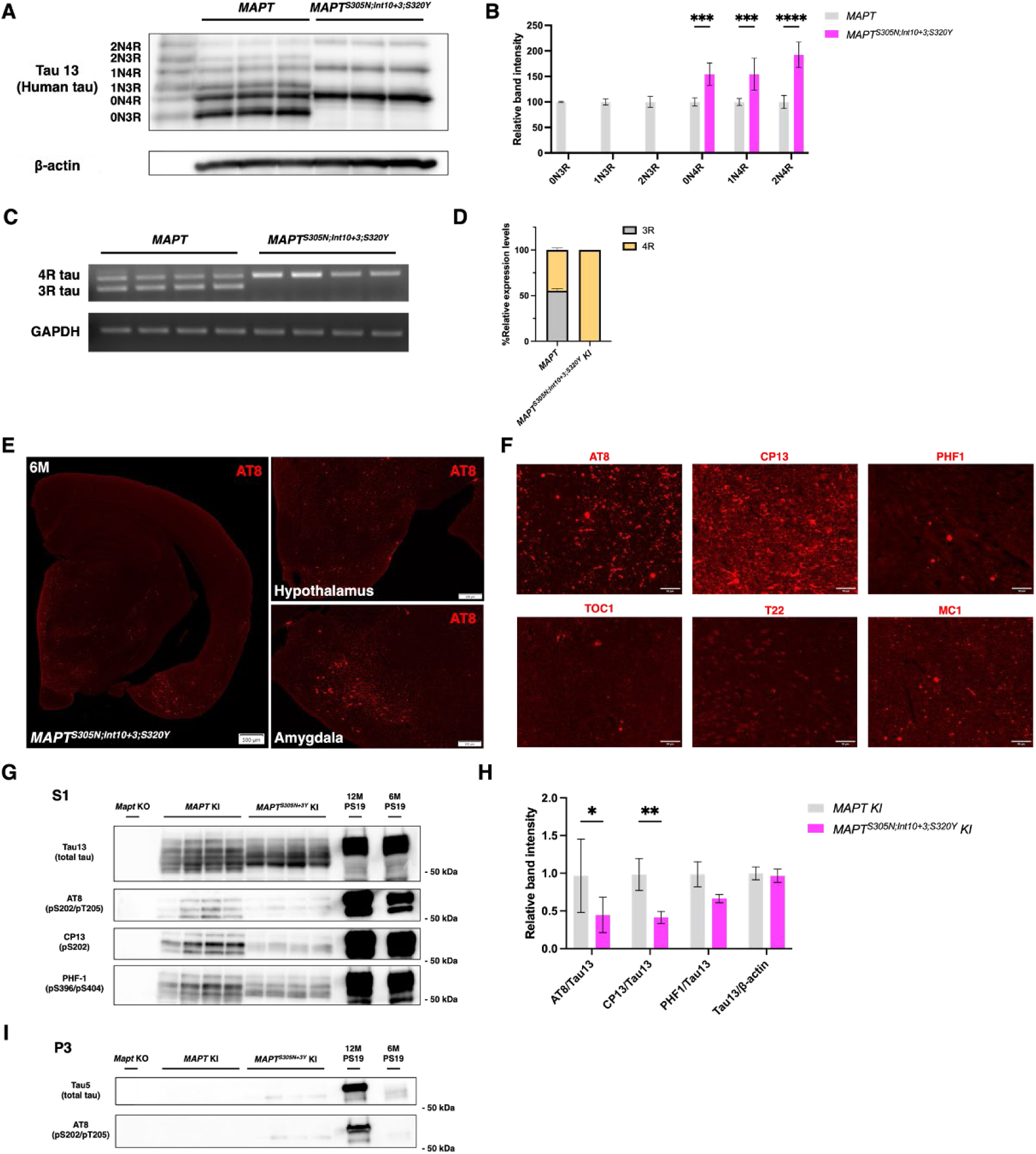
Characterization of *MAPT*^S305N;Int10+3;S320Y^ mice. **A, B.** Western blot analysis of dephosphorylated tau in brain lysates detected by Tau13 antibody (A) and their semi-quantification (B). **C, D.** Reverse transcription PCR analysis of 3R/4R tau (C) and measured ratios (D). **E.** AT8 immunopathology in brain slices from *MAPT*^S305N;Int10+3;S320Y^ mice. **F**. Tau pathology characterization using antibodies described in Figure 3A. **G-H**. Results of sarkosyl fractionation using thalamus of *MAPT*^S305N;Int10+3;S320Y^ mice. Western blot of S1 fraction (G), semi-quantification (H) and Western blotting of P3 fraction (I) are shown.

**Figure S3.**
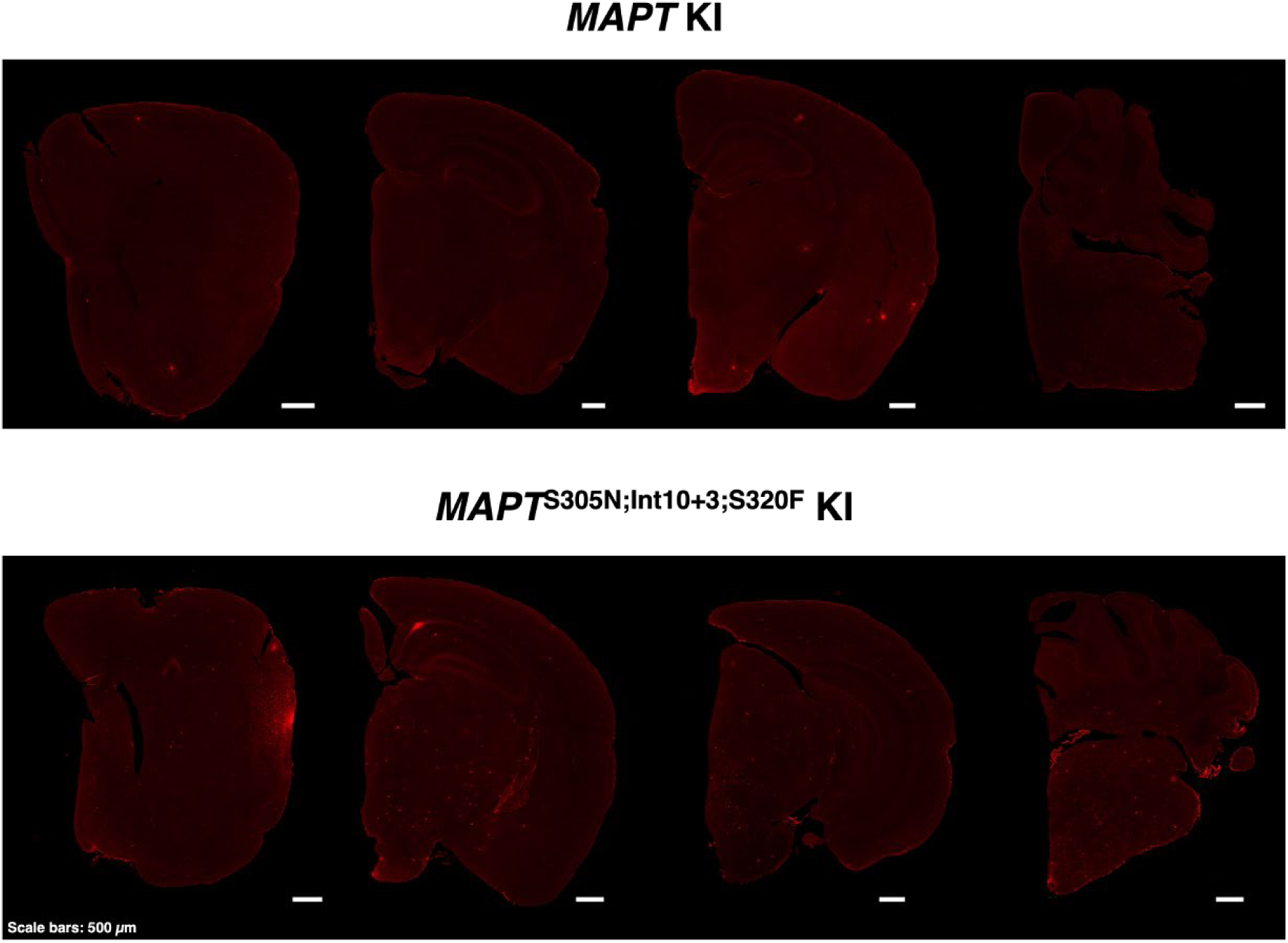
Representative coronal sections of *MAPT*^S305N;Int10+3;S320F^ mice stained with AT8 antibody. These images were used to create heat maps shown in Figure 2C.

**Figure S4.**
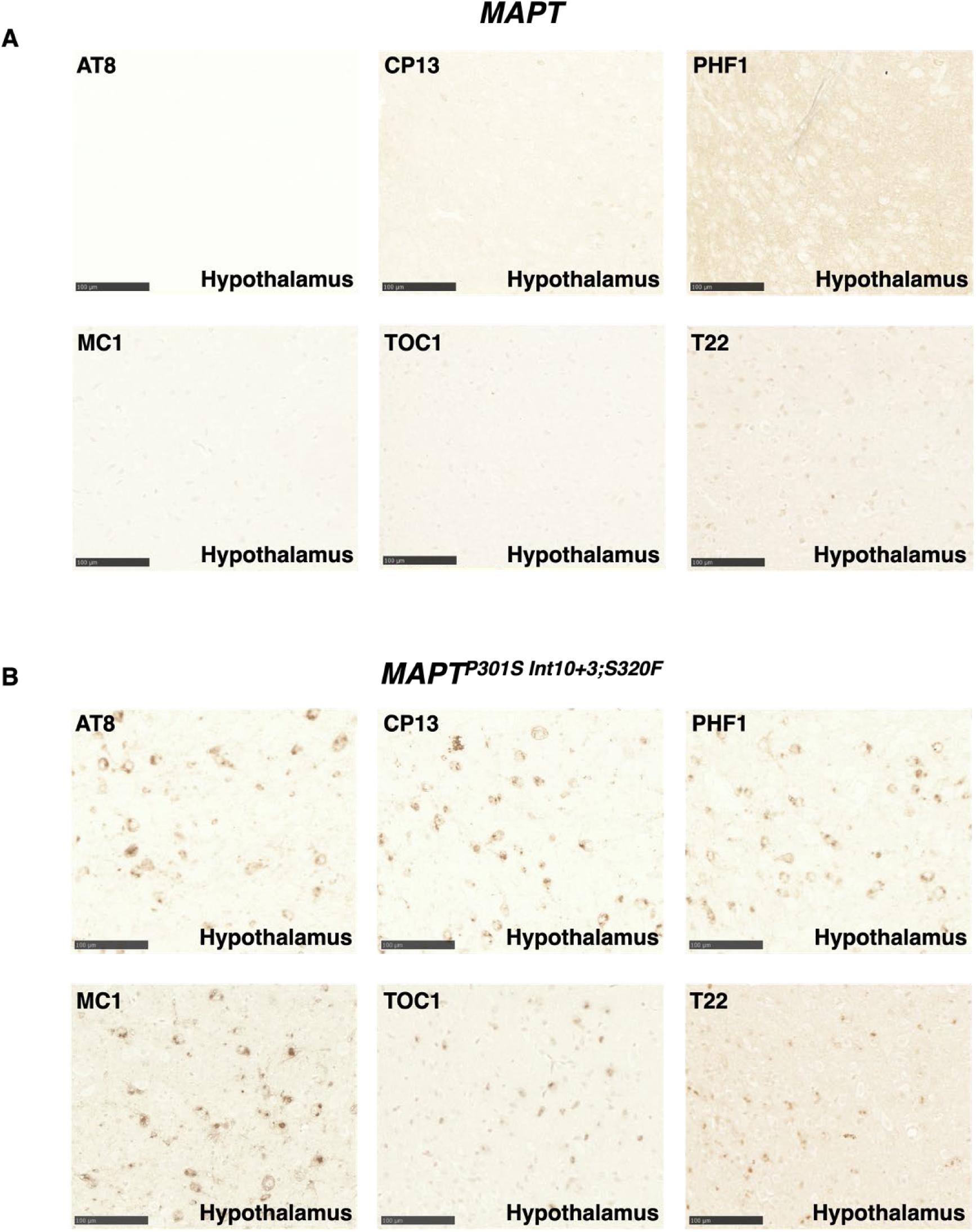
DAB staining of *MAPT^P301S;Int10+3;S320F^* hypothalamus using noted antibodies. Scale bars: 100 µm. Antibodies shown in Figure 3A were used.

**Figure S5.**
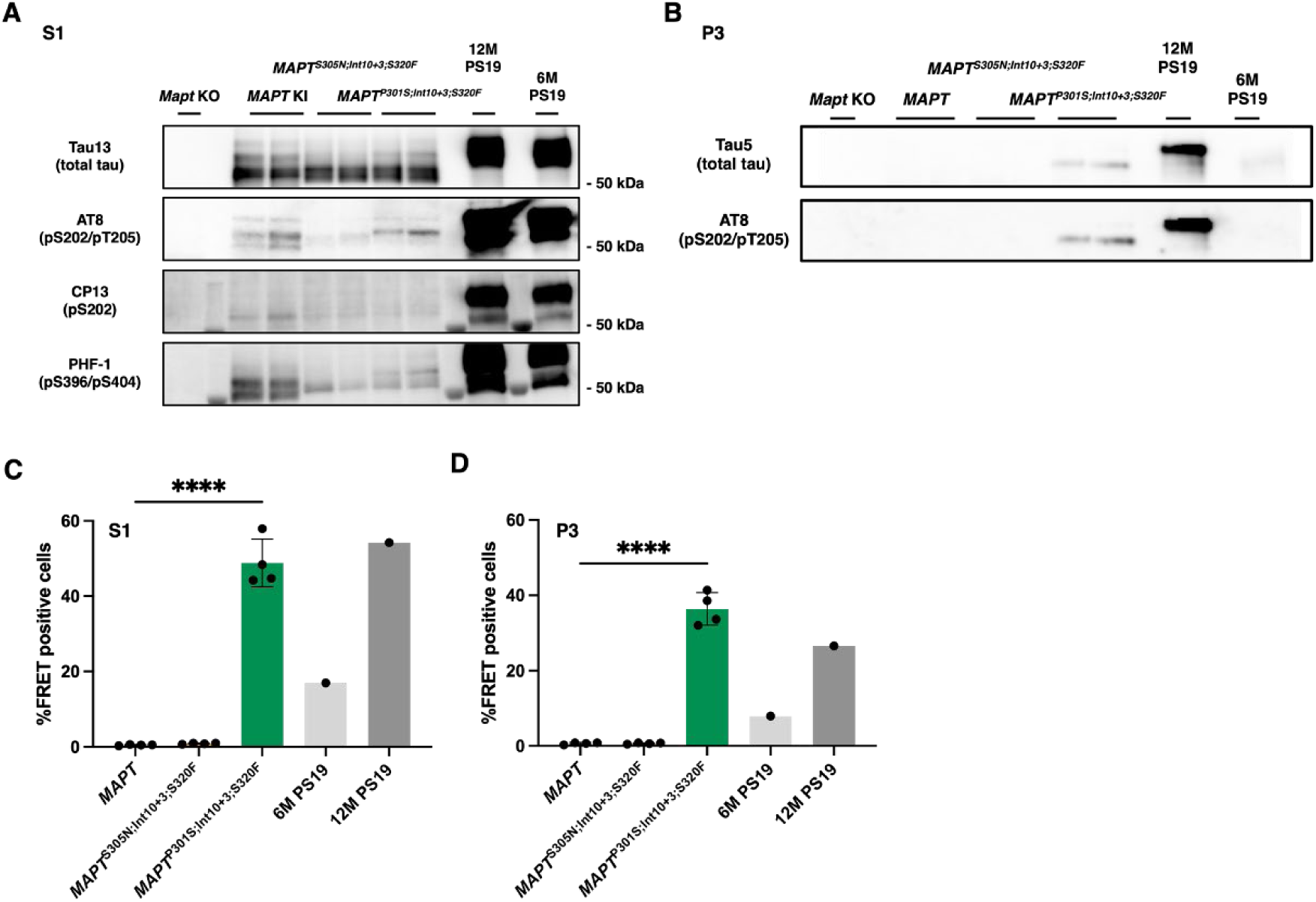
Western blot analysis of tau in S1 and P3 fractions from cortical brain extracts. Brain extracts from mouse cortex of each line were fractioned by sarkosyl fractionation, and obtained S1 (A) or P3 (B) fractions were applied to western blot as described at Figure 4A-D.

**Figure S6.**
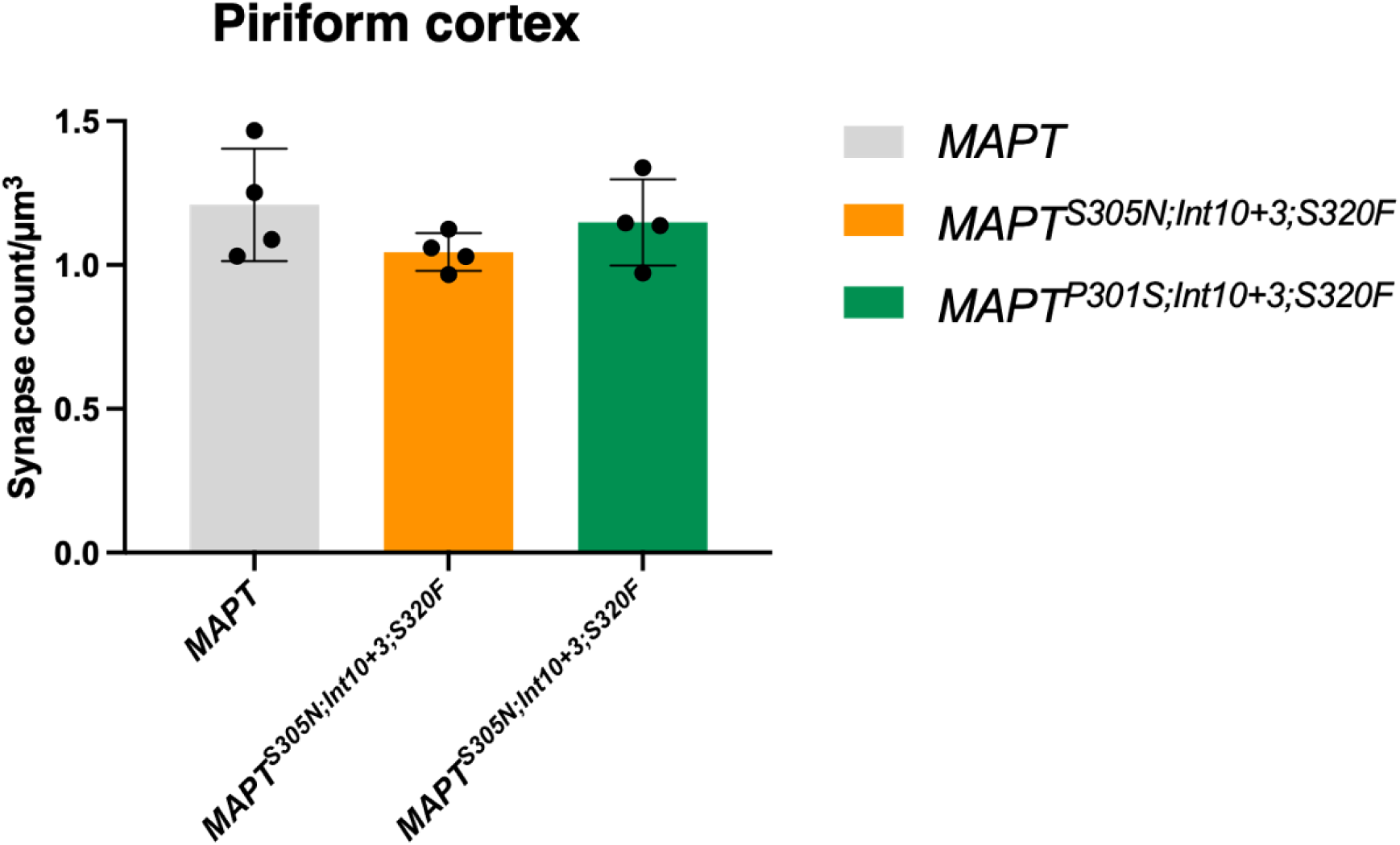
Synapse colocalization analysis of piriform cortex as a negative control of Figure 5D. Synapse counts per volume (µm^3^) of the obtained confocal images are shown.

**Figure S7.**
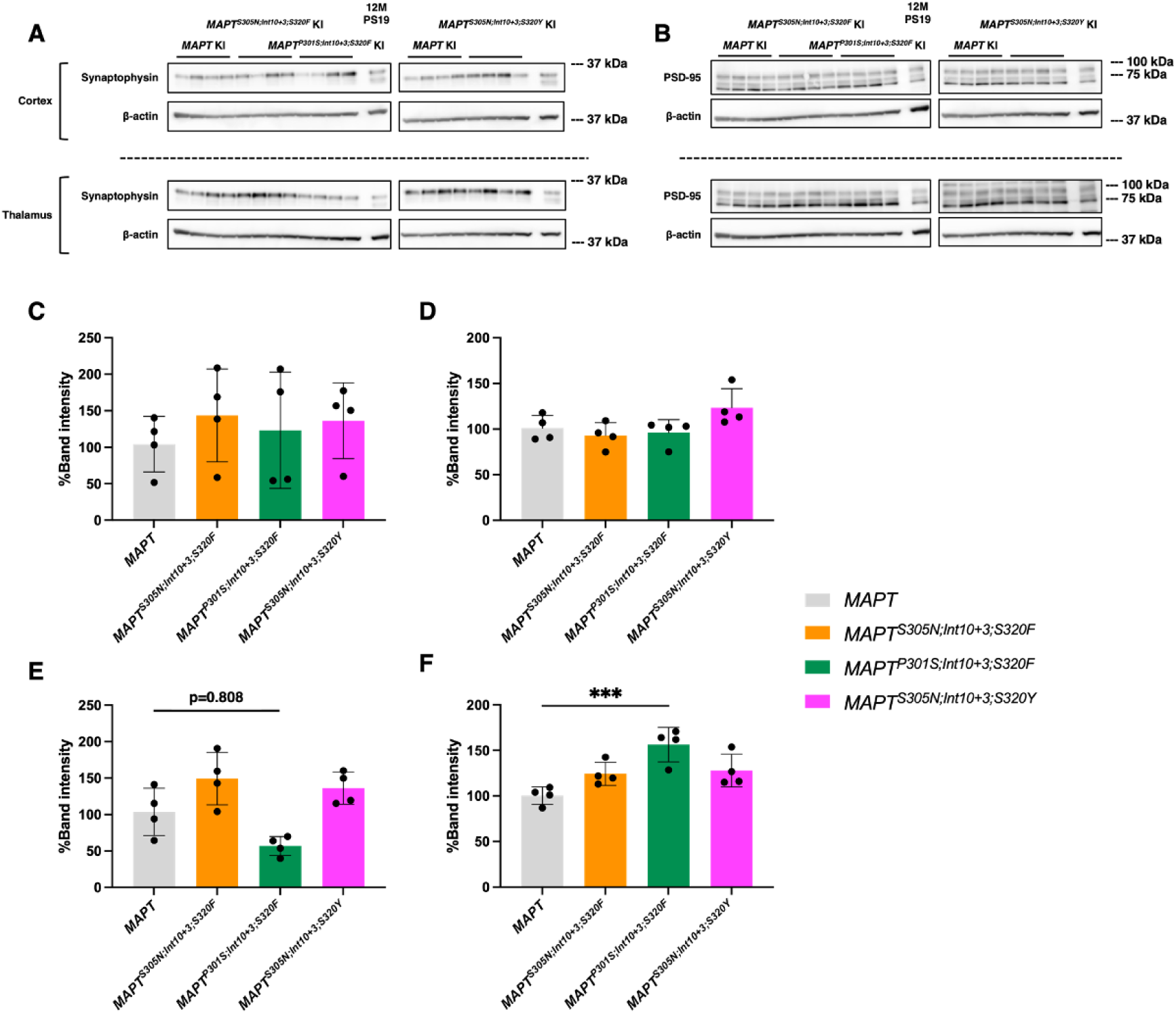
Western blot analysis of pre-synaptic marker (synaptophysin) and post-synaptic marker (PSD-95) in the brain extracts of cortex or thalamus. **A-F.** Western blot analysis of synaptophysin (A) and PSD-95 (B) in cortex (upper) or thalamus (lower) and its semi-quantification (synaptophysin: cortex (C), thalamus (D), PSD-95: cortex (E) and thalamus (F)).

**Figure S8.**
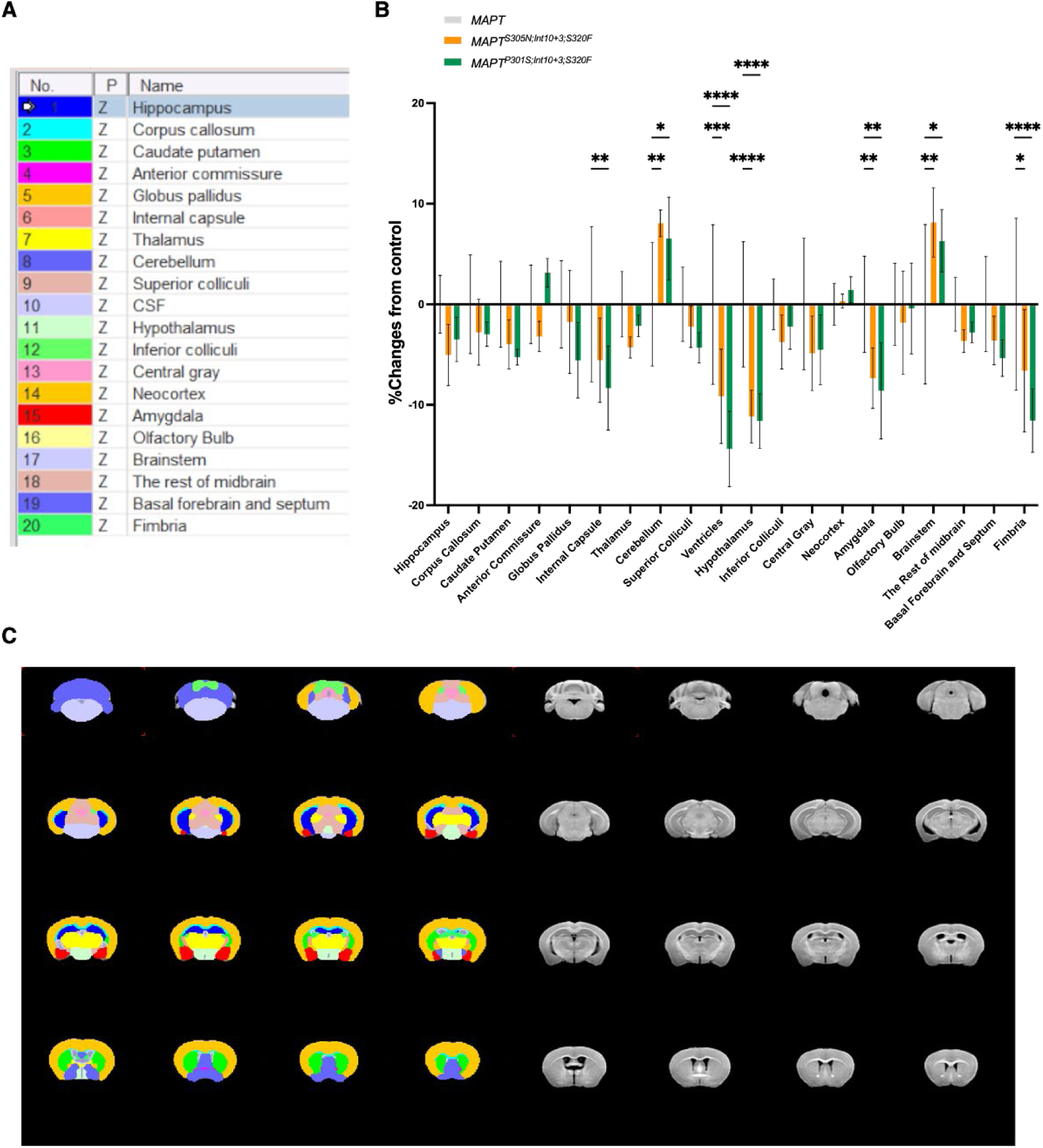
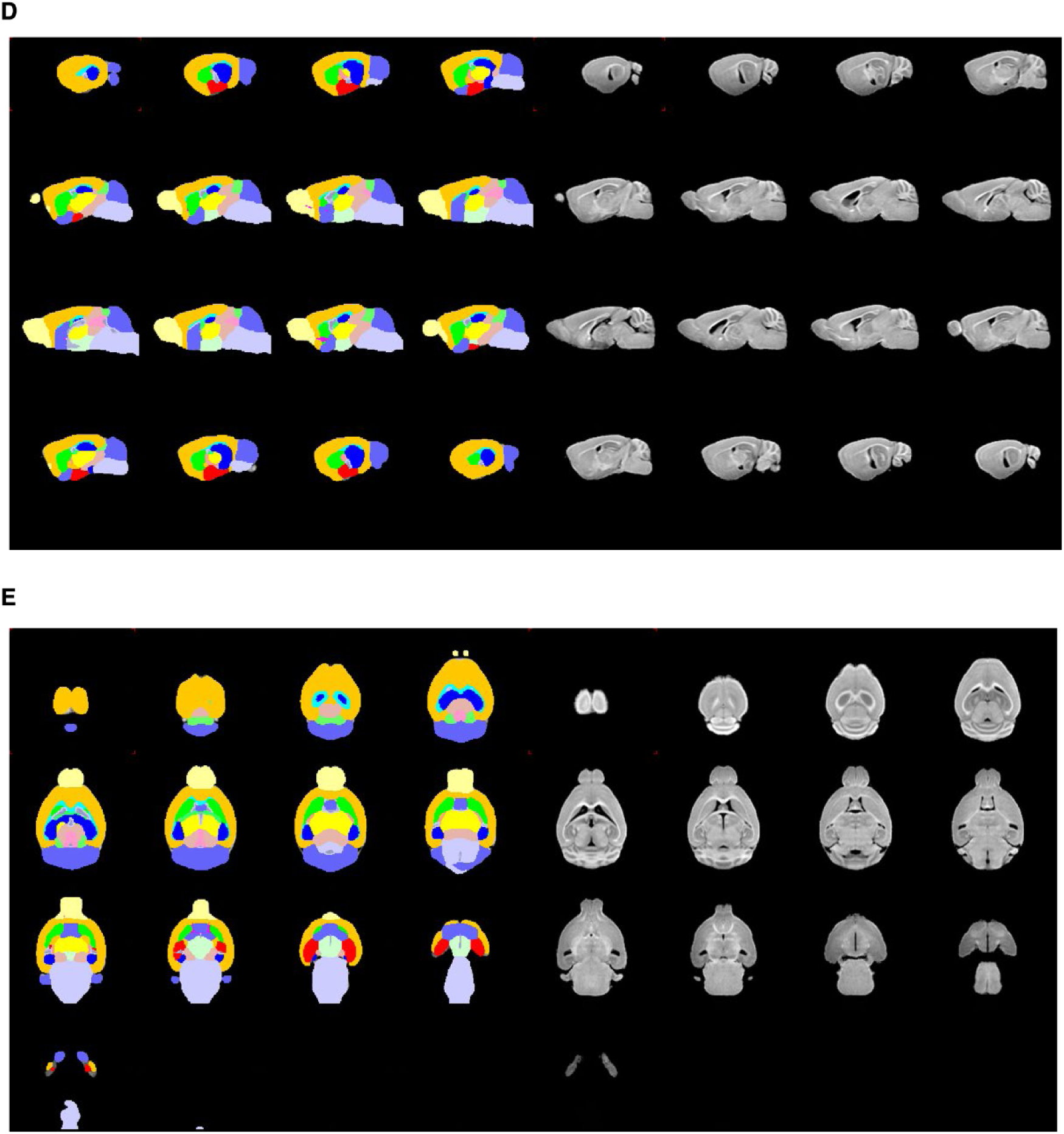
MRI-based volumetric analysis of triple mutant mouse lines. **A.** Selected regions of interest (20 regions) in the mouse brain. **B.** Difference of mutant mice brain volumes in the 20 regions compared with MAPT KI averaged brain regions. **C-E.** Coronal (C), sagittal (D) and transverse (E) overviews (slice pitch, 0.5 mm) of the mouse MR template control averaged brain containing 20 regions colored as indicated in A.

**Figure S9.**
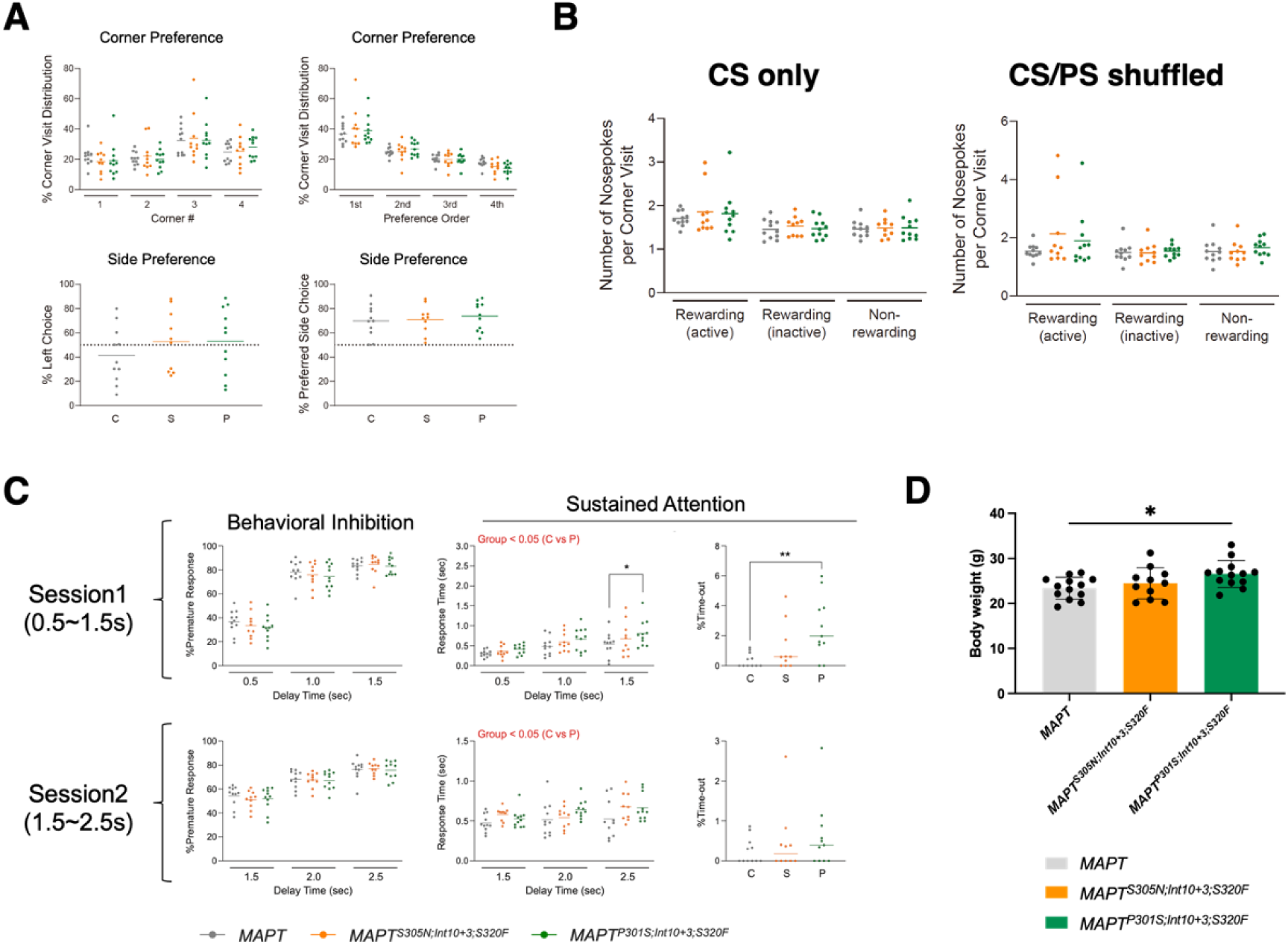
Additional IntelliCage behavioral analysis results not shown at Figure 6. **A.** Corner preference and side preference of three mouse lines during basal activity measurement. **B.** Number of nosepokes per corner visit during learning flexibility tests (CS only: left, CS/PS shuffled: right). C. Data for sessions 1 and 2 behavioral inhibition or sustained attention tests. **D**. Body weight of the three lines of female mice at the end of the serial behavior tests.

### Supplementary Tables

**Table S1.**
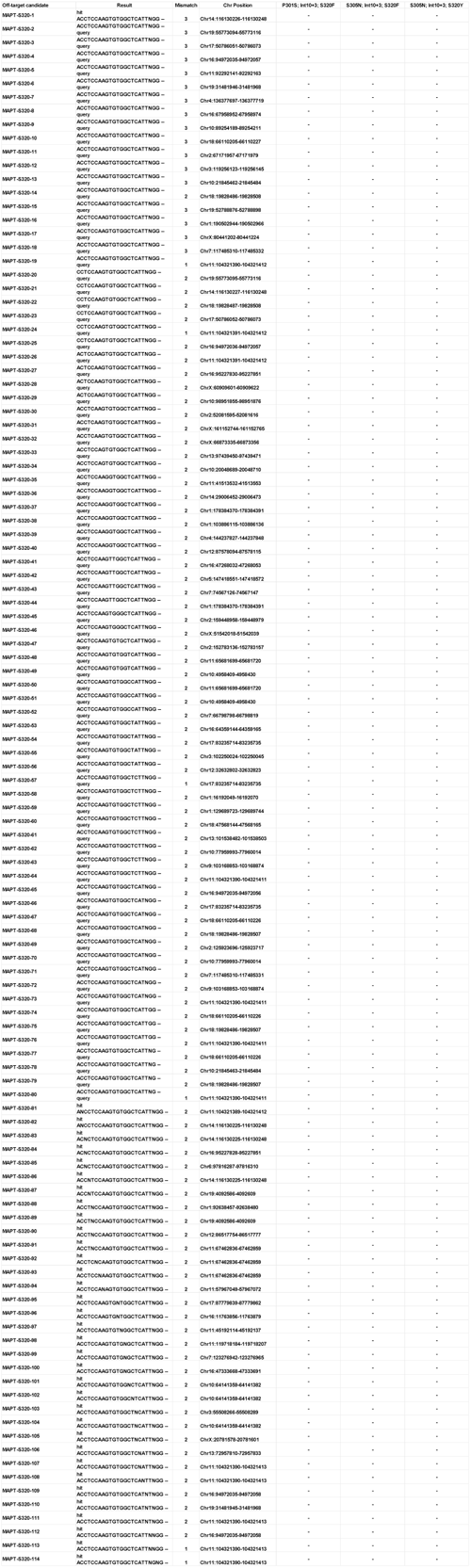
Off-target analyses of the mutant lines generated in the present study. (The original Excel file will be available upon request.)

**Table S2.** Antibodies used in the present study.

| Antibody | Source | Dilutions for WB | Dilutions for IHC |
| --- | --- | --- | --- |
| CP13 | Kindly provided by Peter Davis | 1:1000 | 1:3000 |
| AT8 | Inogenetics #90206 (anti-PHF-TAU) | 1:1000 | 1:200 |
| PHF-1 | Kindly provided by Peter Davis | 1:1000 | 1:500 |
| TOC1 | NOVUS NBP3-20163 |  | 1:1000 |
| T22 | Sigma-Aldrich, ABN454 |  | 1:200 |
| MC1 | Kindly provided by Peter Davis |  | 1:200 |
| Tau13 | Santa Cruz #sc-21796 | 1:2000 or 1:5000 |  |
| Tau5 | Thermo #AHB0042 | 1:1000 |  |
| MAP2 | Abcam ab183830 |  | 1:8000 |
| KIF5A | Proteintech 21186-1-AP |  | 1:100 |
| GA5 (Anti-GFAP) | Merck, MAB360 |  | 1:200 |
| Anti-Iba1 | Fujifilm-wako 013-27691 |  | 1:200 |
| Synaptotagmin | Synaptic System 105-002 |  | 1:500 |
| Homer1 | Synaptic System 160-011 |  | 1:500 |
| A60 (Anti-NeuN) | Merck MAB377 |  | 1:500 |
| Anti-cleaved Caspase-3 | Cell signaling #9661 |  | 1:400 |
| N1D (Anti-Aβ) | Lab-made (Saido et al. 1994) | | 1:200 |
| Anti-β-actin, HRP | proteintech HRP-60008 | 1:30000 | |
| Anti-Rabbit IgG, HRP | proteintech SA00001-2 | 1:5000 |  |
| Anti-Mouse IgG, HRP | proteintech SA00001-1 | 1:5000 or 1:10000 |  |

